# Mapping the Fragment-Binding Landscape of SARS-CoV-2 nsp14 for Antiviral Drug Discovery

**DOI:** 10.64898/2026.10.21.760455

**Authors:** Xinyue Shi, Qian Wang, Danni Dong, Manming Xu, Fatma Al Busaidi, Matthew W. Bowler, Shozeb Haider, Frank Kozielski

## Abstract

Non-structural protein 14 (nsp14) is a bifunctional coronavirus drug target, combining a 3’-to-5’ exoribonuclease domain with an N7-methyltransferase domain. Proof-of-concept data have demonstrated beneficial effects in animal models upon inhibition. Using microscale thermophoresis and X-ray crystallography, we screened full-length nsp14 for fragment hits and developed SAR-by-catalogue analogues with five-to eight-fold improved affinity. Several scaffolds showed pronounced overlap between distinct pockets, complicating structure-based optimisation. Analysis of mutational frequencies across over 14 million SARS-CoV-2 genome sequences identified nsp14 residues with markedly high and very low mutation rates, highlighting sites best suited to pan-coronavirus inhibitor design. Combining these data with all 19 publicly known fragment- and ligand-binding regions on nsp14, most of which have never been directly compared, we introduce a systematic site-naming system and rank every site by its potential for future structure-based development. Weighing criteria include proof-of-principle inhibitor data, cross-coronavirus conservation, mutation frequency, the number of fragments bound per site, and susceptibility to crystallographic artefacts from symmetry mates. This ranked, unified binding-site atlas resolves which of the many previously reported nsp14 sites merit prioritisation, providing a practical resource for future coronavirus antiviral drug development.

## Introduction

More than 779 million confirmed Coronavirus Disease 2019 (COVID-19) cases have been recorded worldwide by the World Health Organization (WHO) since the start of the pandemic (World Health Organization, 2026). COVID-19 is caused by a positive-sense single-stranded RNA (+ssRNA) virus named Severe Acute Respiratory Syndrome Coronavirus 2 (SARS-CoV-2).

SARS-CoV-2 consists of four structural proteins, sixteen non-structural proteins and six accessory proteins (Mariano *et al*., 2020; Briggs *et al*., 2023). Non-structural proteins are important in the synthesis of negative-sense single-stranded RNA (-ssRNA), the replication of +ssRNA and the transcription of -ssRNA to subgenomic messenger RNA. One of the key proteins in the Replication Transcription Complex (RTC) is nsp14, a bifunctional protein, which contains both 3’-5’ exoribonuclease (ExoN) and N7-methyltransferase (N7-MTase) activities (Minskaia *et al*., 2006; Chen *et al*., 2009). The ExoN domain can remove mismatched nucleotides from the 3’-end of RNA following viral RNA replication (Minskaia *et al*., 2006; Bouvet *et al*., 2012). The activity of the nsp14 ExoN domain has been shown to be enhanced by the binding of nsp10 (Hsu *et al*., 2021). The N7-MTase domain transfers the methyl group from its substrate S-adenosyl-L-methionine (SAM) to the 7^th^ position of guanine of viral mRNA (Chen *et al*., 2009; Ghosh and Lima, 2010), generating the by-product S-adenosyl-L-homocysteine (SAH) during the reaction. The viral methylated RNA cap can increase the stability of the mRNA as well as the chances of the virus of escaping from the host immune system (Kakavandi *et al*., 2023). Nsp14 is considered a potential drug target, as its inhibition disrupts virus replication (Subissi *et al*., 2014) and ExoN active site mutants result in non-viable virus (Ogando *et al*., 2020). N7-MTase mutants have also been shown to affect nsp14 translational inhibition activity (Hsu *et al*., 2021).

The understanding of nsp14 function was further advanced by the determination of its structure. Various SARS-CoV-2 nsp14 crystal structures and cryo-electron microscopy reconstructions are available including unbound nsp14 (Czarna *et al*., 2022; Imprachim *et al*., 2023), the nsp14 ExoN domain in complex with nsp10 (Lin *et al*., 2021; Moeller *et al*., 2022), the nsp14 N7-MTase domain (Kottur *et al*., 2023), a nsp10-nsp14-RNA ternary complex (Liu *et al*., 2021) and nsp14 as part of the RTC (Yan *et al*., 2021).

To develop nsp14 inhibitors one of the strategies is to disrupt the binding between nsp14 and nsp10 by targeting the binding interface (Arabi-Jeshvaghani *et al*., 2023). As SARS and SARS-CoV-2 N7-MTase domains do not contain the canonical Rossman fold, the N7-MTase was selectively targeted with inhibitors (Ferron *et al*., 2017; Czarna *et al*., 2022). Several bisubstrate inhibitors, targeting both SAM and RNA binding sites, were designed and developed (Ahmed-Belkacem *et al*., 2022; Jung *et al*., 2022; Georgiou *et al*., 2026). A total of 72 fragment hits has been identified using fragment-based screening by X-ray crystallography (Imprachim *et al*., 2023). Fluorescence-based assays (Basu *et al*., 2021; Bobiļeva *et al*., 2021; Kasprzyk *et al*., 2021; Rona *et al*., 2022; Coker *et al*., 2026) and mass spectrometry (Pearson *et al*., 2021) have also been employed to identify inhibitors.

In this project, nsp14, and its individual ExoN and N7-MTase domains, were screened for fragment hits using microscale thermophoresis (MST). Subsequently, the initial hits were subjected to X-ray crystallography and further investigated with orthogonal biophysical techniques to obtain estimates of their K_d_ values. An initial Structure-Activity Relationship (SAR)-by-catalogue study was conducted by testing commercially available fragment hit analogues for binding. Here we report the identification and characterisation of novel fragment hits targeting nsp14. An initial SAR-by-catalogue study using X-ray crystallography allowed us to conduct a detailed analysis of binding modes. Through a combination of various criteria, all currently known binding sites on both functional domains of nsp14 were ranked for their potential as real binding sites.

## Material and Methods

Various nsp14 constructs were expressed and purified for thermal shift assays (TSA), MST and structural studies followed by fragment soaking.

### Purification of nsp14 constructs for microscale thermophoresis

For fragment screening using MST binding assays as well as the estimation of K_d_ values, three nsp14 constructs containing histags were purified. SUMO was also purified and served as a control.

Full-length nsp14 (nsp14_1-527_) cDNA was subcloned into ppSUMO-2 resulting in an expression vector coding for an N-terminal histag and a SUMO fusion protein, an Ulp1 cleavage site and, for *E. coli* expression, codon-optimised cDNA coding for nsp14. The resulting plasmid was transformed into Tuner (DE3) *E. coli* cells and incubated on an agar plate supplemented with 50 μg/ml kanamycin at 37°C overnight. A single colony was picked from the plate and added into 30 ml TB medium supplemented with 50 μg/ml kanamycin. The small culture was then placed in a shaker-incubator at 37°C overnight. Then, the small culture was separately added into six 2 L Erlenmeyer flasks each containing 1 L TB medium supplemented with 50 μg/ml kanamycin. The cultures were placed in a shaker-incubator at 37°C until their OD_600_ reached 0.8. The cultures were then induced with 0.1 mM IPTG and incubated at 20°C for 24 h. The culture was harvested by centrifugation at 4000 rpm (Avanti® J-E centrifuge from Beckman Coulter, rotor: JLA 16.250) for 20 min at 4°C. The cell pellet was resuspended with 200 ml 50 mM Tris-HCl pH 8.4, 300 mM NaCl and 10 mM imidazole, frozen in liquid nitrogen and stored at −80°C.

For purification, the cell pellet was thawed and sonicated for ten rounds on ice. The lysate was centrifuged for 90 min at 20000 rpm, 4°C (Avanti® J-E centrifuge from Beckman Coulter, rotor: JLA 25.50). The supernatant was collected and loaded into a 5 mL HisTrap FF crude column (Cytiva) pre-equilibrated with buffer A (50 mM Tris-HCl pH 7.8, 300 mM NaCl, 10 mM Imidazole and 1 mM PMSF). The column was then washed with 50 column volumes (CVs) of buffer B (50 mM Tris-HCl pH 7.8, 300 mM NaCl and 30 mM Imidazole). Protein bound to Ni-NTA was eluted with buffer C (50 mM Tris-HCl pH 7.8, 500 mM NaCl and 500 mM Imidazole), collected in 2 ml fractions and the fractions were analysed by SDS-PAGE. The fractions containing nsp14 were combined and quantified using the Bradford reagent.

The nsp14 ExoN domain (ExoN_1-289_) and the N7-MTase domain (nsp14_283-527_) were expressed and purified following methods described previously (Kozielski *et al*., 2022). SUMO was a purification by-product when cleaved off by Ulp1 protease before the second Ni-NTA column as it retains its histag and binds to Ni-NTA. It was subsequently eluted with imidazole to yield pure SUMO for subsequent control measurements.

### Expression and purification of full-length nsp14 for structural studies

Nsp14 was expressed, purified and crystallised as previously described (Imprachim *et al*., 2023), with minor modifications. The nsp14_7-527_ cDNA was subcloned into expression vector ppSUMO-2. BL21-Gold(DE3) cells (Agilent Technologies) and TB modified medium (Melford) with 50 µg/ml Kanamycin were used for expressing nsp14_7-527_ at 37°C. When the OD_600_ reached 0.6-1.0, IPTG was added to the culture to a final concentration of 1 mM. After incubating at 220 rpm at 18°C for 20 to 24 h, the culture was centrifuged at 4°C at 8000 g for 20 min. Cell pellets were resuspended in lysis buffer (50 mM HEPES pH 8.0, 500 mM NaCl, 10 mM Imidazole, 5% Glycerol, 1 mM TCEP and 1 mM PMSF) and flash frozen in liquid nitrogen.

The cell suspension was thawed and sonicated on ice 14 times for 30 sec to break the cells. Between each sonication cycle, the cell suspension was placed for 1 min on ice. The sample was then centrifuged at 4°C at 20000 rpm for 1 h. The supernatant was loaded into a 5 ml HisTrap FF column (Cytiva) previously equilibrated with lysis buffer. The column was connected to a Biorad FPLC and washed with 50 CVs of wash buffer (50 mM HEPES pH 8.0, 500 mM NaCl, 20 mM Imidazole, 5% Glycerol and 1 mM TCEP), followed by 15 CVs of 1^st^ column elution buffer (50 mM HEPES pH 8.0, 500 mM NaCl, 300 mM Imidazole, 5% Glycerol and 1 mM TCEP). Samples showing an elution peak were tested using SDS-PAGE. The samples containing nsp14 were added into a 10K MWCO SnakeSkin^TM^ dialysis tubing with 1 mg Ulp1 for every 30 mg nsp14 and placed in 2 L dialysis buffer (50 mM HEPES pH 7.5, 500 mM NaCl, 2 mM DTT and 5% Glycerol) for 18 h. The sample was then loaded into a fresh HisTrap FF column. Cleaved nsp14 was eluted with six CVs of 2^nd^ column elution buffer (50 mM HEPES pH 7.5, 500 mM NaCl, 30 mM Imidazole, 5% Glycerol and 1 mM TCEP). Gel filtration was then performed with the eluted sample as described (Imprachim *et al*., 2023). Fractions containing nsp14 were concentrated to 10 mg/ml for subsequent crystallisation and TSA experiments.

### Purification of nsp14 constructs for thermal shift assays

For TSA experiments, nsp14 constructs were purified by two affinity purification steps to remove the SUMO and histidine tags.

The nsp14 ExoN domain was expressed and purified as previously described (Kozielski *et al*., 2022). ExoN was then pooled and cleaved overnight with Ulp1 protease supplemented with 3 mM DTT. Simultaneously, the protein was also dialysed against buffer containing 50 mM Tris-HCl pH 7.5 and 300 mM NaCl. Thereafter, the protein was passed through the Ni-NTA column for a second time and was eluted with 50 mM Tris-HCl pH 7.5, 300 mM NaCl and 20 mM imidazole. The cleaved protein, without the His-SUMO fusion tag, was captured in the flow-through, which was then pooled and dialysed for 18 h in a buffer containing 50 mM Tris-HCl pH 7.5 and 300 mM NaCl. The protein was then concentrated to 2 mg/mL using an Amicon ultraconcentration device (Millipore).

The N7-MTase domain (nsp14_289-525_) was expressed and purified as described above for nsp14_7-527_ using BL21-Rosetta2 (DE3) cells. The following buffers were used during purification. Lysis buffer (50 mM Tris-HCl pH 7.8, 300 mM NaCl, 10 mM imidazole and 1 mM PMSF), 1^st^ column wash buffer (50 mM Tris-HCl pH 7.8, 300 mM NaCl and 20 mM imidazole), 1^st^ column elution buffer (50 mM Tris-HCl pH 7.8, 300 mM NaCl and 200 mM imidazole), Dialysis buffer (60 mM HEPES pH 8.0, 500 mM NaCl and 3 mM DTT), 2^nd^ column elution buffer (60 mM HEPES pH 8.0, 500 mM NaCl and 35 mM imidazole) and final buffer (60 mM HEPES pH 8.0, 500 mM NaCl).

### Microscale thermophoresis assays

To label nsp14 constructs for MST measurements, SUMO-nsp14_1-527_, SUMO-nsp14_1-289_ and SUMO-nsp14_283-527_ were diluted in MST assay buffer to 800 nM. The MST assay buffer for nsp14_1-527_ and nsp14_1-289_ was 50 mM NaPO_4_ pH 7.5, 150 mM NaCl and 0.05% Tween-20, and for nsp14_283-527_ was 60 mM HEPES pH 7.4, 150 mM NaCl and 0.1 % PEG-8000. The Red-Tris-NTA 2^nd^ generation dye was diluted to 100 nM. 50 μl nsp14 constructs were incubated with 50 μl dye on ice for 1 h and centrifuged at 4°C at 13000 rpm for 5 min.

One hundred fragments were tested for binding to nsp14 constructs using the “binding check” procedure. Labelled nsp14 constructs were further diluted to 20 nM in MST assay buffer. Fragment stocks (200 mM) were diluted with MST assay buffer to either 20 mM or 10 mM. DMSO was diluted with MST assay buffer with the same 10-fold or 20-fold ratio as the fragment stock. For the target-only sample, diluted nsp14 constructs were mixed with the same volume of diluted DMSO. For the complex sample, diluted nsp14 constructs were mixed with the same volume of 20 mM or 10 mM fragments. After 30 min incubation at room temperature, the samples were then loaded into Monolith standard capillaries and measured by the Monolith NT.115 instrument. The measurements were performed under 5 - 15 sec MST on time, 100% excitation power and 40% MST power. The data was analysed using MO.Control and MO.Affinity Analysis software.

Nsp14_7-527_ labelled with RED-NHS 2^nd^ generation dye was used to measure the binding affinity between nsp14 constructs and fragments. Protein labelling followed the instructions in the NanoTemper Protein Labelling Kit RED-NHS 2nd Generation manual. Specifically, 100 µL 10 µM nsp14_7-527_ was buffer exchanged with the A-column in the kit to labelling buffer NHS (130 mM NaHCO_3_ pH 8.2-8.3 and 50 mM NaCl). 90 µL 10 µM nsp14_7-527_ was then mixed with 10 µL 300 µM RED-NHS dye. After incubating for 30 min at RT in the dark, the mixture was loaded onto the B-column provided in the kit. 550 µL PBS-T (Phosphate-buffered saline with 0.1% Tween-20) buffer was added to the column to remove the free dye. The labelled nsp14_7-527_ was then eluted by adding 450 µL PBS-T buffer. For cysteine-directed labelling of nsp14_7-527_, the NanoTemper Protein Labelling Kit RED-MALEIMIDE 2nd Generation was used according to the manufacturer’s protocol.

20 nM labelled nsp14 constructs were mixed with each fragment in 16 wells in 384-well microplates. Each ligand was diluted to final concentrations of 10 mM to 305 nM by a two-fold dilution series. The final concentration of nsp14 constructs in each well was 10 nM. The mixed samples were incubated at room temperature for 1 h and loaded into 16 Monolith NT.115 Premium capillaries and measured by the Monolith NT.115 instrument. The measurements were performed under 5 - 15 sec MST on time, 40% MST power, 100% excitation power for nsp14_283-527_ and 30% excitation power for nsp14_7-527_.

### Thermal shift assays. Preparation of nsp14

As a DMSO control, 3.5 *µ*L of 14 mg/mL nsp14 and 1 *µ*L of DMSO were mixed with 95.5 *µ*L buffer A (50 mM HEPES pH 7.5, 500 mM NaCl, 5% glycerol and 1 mM TCEP). 10 *µ*L of 0.5 mg/mL nsp14 with 1% DMSO were loaded into Tycho capillaries and measured in triplicate. For subsequent measurements in the presence of fragments, 3.5 *µ*L of 14 mg/mL nsp14, 1 *µ*L of 200 mM fragment, and 95.5 *µ*L buffer A were mixed and loaded into Tycho capillaries and measured in triplicate.

### Preparations of the ExoN domain (nsp14_1-289_)

As a control, 10 *µ*L of 5 mg/mL nsp14_1-289_, 1 *µ*L of DMSO and 89 *µ*L buffer B (50 mM HEPES pH 7.0 and 300 mM NaCl) were mixed. 10 *µ*L of 0.5 mg/mL nsp14_1-289_ with 1% DMSO were loaded into Tycho capillaries and measured in triplicate. For measurements of fragments, 10 *µ*L of 5 mg/mL nsp14_1-289_ and 1 *µ*L of 200 mM fragment were mixed with 89 *µ*L buffer B and loaded into Tycho capillaries and measured in triplicate.

### Dilution of the N7-MTase domain (nsp14_289-525_)

5 *µ*L of 10 mg/mL nsp14_289-525_, 1 *µ*L of DMSO and 94 *µ*L buffer C (60 mM HEPES pH 8.0 and 500 mM NaCl) were mixed as control. 10 *µ*L of 0.5 mg/mL nsp14_289-525_ with 1% DMSO were loaded into Tycho capillaries and measured in triplicate. 5 *µ*L of 10 mg/mL nsp14_289-525_, 1 *µ*L of 200 mM fragment, and 94 *µ*L buffer C were mixed and loaded into Tycho capillaries and measured in triplicate. The heating temperature increased from 35°C to 95°C in 3 min. The inflection temperature (T_i_) of the proteins was automatically calculated by the Tycho NT.6 instrument following the heating process.

### Crystallisation and fragment soaking

Crystallisation drops were set up using the reported condition of 1.26 M sodium phosphate monobasic (NaH_2_PO_4_) and 0.14 M potassium phosphate dibasic (K_2_HPO_4_) using the hanging drop vapour diffusion method. Crystals appeared after 30 days at 4°C. To reduce crystal growth time and increase the crystal size, seed stock 1 was prepared with crystals from the initial drops. Seed beads (Hampton Research) were used following the reported protocol (Hampton Research, 2024), 1 µl of 5 mg/ml nsp14 was mixed with 1 µl seed stock 1. Rod clusters appeared five days after setting up drops at 18°C. These new crystals were used to prepare seed stock 2 and diluted to a 1:10 ratio resulting in seed stock 3 to reduce the number of seeds. New drops were set up by mixing 2 µl of 5 mg/ml nsp14 with 2 µl seed stock 3 and incubated at 18°C. Large single crystals were observed seven days after setting up drops. The reservoir in all wells contains 500 µl 1.26 M NaH_2_PO_4_ and 0.14 M K_2_HPO_4_. Crystals were soaked with solutions containing 1.5 M NaH_2_PO_4_, 0.17 M K_2_HPO_4_ and 50 mM fragments dissolved in DMSO for 1 h, then harvested with cryo-loops and flash frozen in liquid nitrogen.

### Data collection, structure determination and refinement

Fragment data were collected at the European Synchrotron Radiation Facility (ESRF), beamlines ID23-1 and ID23-2. Analogue data were collected at ESRF beamline ID30A-1 MASSIF-1 (Bowler *et al*., 2015) and Diamond Light Source (DLS) beamline I03 using unattended data collection. X-ray centring was used to locate the optimal crystal region for data collection (Svensson *et al*., 2015). Images (980 to 2300) were collected with either 0.1° or 0.05° oscillation and 0.01 or 0.02 sec exposure time.

Diffraction data were automatically integrated and scaled by autoPROC (Vonrhein *et al*., 2011), EDNA Autoprocessing, GrenADES parallelproc (Monaco *et al*., 2013) and XIA2_DIALS (Winter, 2010; Winn *et al*., 2011; Winter *et al*., 2018). Molecular replacement was performed using Phaser-MR (McCoy *et al*., 2007) from the PHENIX suite (version:1.19.2-4158-000) (Liebschner *et al*., 2019) and SARS-CoV-2 nsp14 (PDB entry 7QGI) was used as the search model. The models were manually refined in Coot (0.9.8.6) (Emsley *et al*., 2010) while inspecting electron density and difference density maps. The structures were then refined using phenix.refine (Afonine *et al*., 2012).

### Calculation of Ro3 and initial physicochemical parameters

MW, ClogP and tPSA were obtained from Chemdraw, version 20.0.038 whereas the number of hydrogen bond acceptors and donors as well as the calculated solubility was obtained using the Molsoft server (https://molsoft.com/mprop/mprop.cgi). Nsp14-ligand interactions were analysed using PLIP (Protein Ligand Interaction Profiler) using the default thresholds (Schake *et al*., 2025). Figures showing protein structures were generated using Open-Source PyMOL (version 2.5.0). The chemical structures of small molecules were generated in Chemical Sketch Tool (https://www.rcsb.org/chemical-sketch).

### Sequence Selection and alignment

Global Initiative on Sharing All Influenza Data (GISAID) is a global online database that provides genomic data for influenza viruses and coronaviruses, including SARS-CoV-2 (https://www.gisaid.org) (Khare *et al*., 2021). In this study, the whole genome sequence of all uploaded SARS-CoV-2 strains was downloaded in FASTA format. The nsp14 sequences were then extracted from the dataset, and any sequences containing unknown residues (represented as ‘X’) were removed. After this filtering process, 14,246,785 NSP14 sequences remained for further analysis.

To improve computational efficiency, we did not conduct a global multiple sequence alignment across all 14 million sequences. Instead, each sequence was individually aligned to the reference Wuhan/WIV04/2019 genome using Parasail (Daily, 2016), allowing for a more effective identification and quantification of mutations. In-house Python scripts were used to process the large dataset, and the mutation rate of each mutation was calculated as the total occurrence of the mutation divided by the total number of sequences. The compiled mutation data were then stored in an Excel sheet.

### Ranking of all known potential of fragment binding sites for subsequent inhibitor development

The main criterion for ranking all described binding sites was the existence of proof-of-concept data for nsp14-targeting compounds developed by structure-based design. The second criterion was the potential impact a fragment binding to a particular site, evaluated by combining available structural information about nsp14 and its binding partners. These include the nsp14-SAH complex (PDB entry 7R2V), the nsp14-TDI-015051 complex (PDB entry 8R7B), the RTC (7EGQ), nsp14 bound with 7MeGpppG (PDB entry 7QIF) and a ternary nsp14-nsp10-RNA complex (PDB entry 7N0B). A third criterion was to assess if fragment binding sites in the nsp14 crystal are formed by nsp14 and one of its symmetry mates. We individually and visually inspected all binding sites by generating nsp14 and its symmetry mates in Coot. The availability of binding data was also evaluated. Estimates of K_d_ values from orthogonal biophysical assays for sites formed by nsp14 and a symmetry mate would indicate that these are still valid binding sites. A further criterion was the number of fragments bound per site, as many fragments binding to the same site would allow for future fragment merging and linking. In contrast, a single fragment bound to a particular site would only allow fragment growing, reducing its potential for future development. We also ranked which of the 19 binding sites on nsp14 are the most conserved between SARS-CoV, MERS-CoV and SARS-CoV-2 coronaviruses with potential to develop pan-nsp14 inhibitors. Therefore, we carried out protein sequence alignments for these three medically relevant CoVs and calculated the conservation of the 19 binding sites. Finally, to assess which of the published nsp14-targeting fragment hits are potential Pan-Assay Interference Compounds (PAINS), the chemical structures of the fragments were extracted from the PDB entries and submitted to the following two PAINS servers: http://www.cbligand.org/PAINS/ (Baell and Holloway, 2010); https://mobyle2.rpbs.univ-paris-diderot.fr/cgi-bin/portal.py#forms::FAF-Drugs4 (Lagorce et al., 2015).

## Results

SARS-CoV-2 nsp14 consists of two major domains connected via a central hinge region (Fig 1A). The ExoN domain is located at the N-terminus whereas the N7-MTase domain is positioned at its C-terminus. Its binding partner, the scaffolding protein nsp10, attaches to nsp14 through the ExoN domain (Fig 1B). To carry out biophysical and structural assays and experiments, a variety of nsp14 constructs covering nsp14 and its individual functional domains, ExoN and N7-MTase, were expressed and purified (Fig 1A). For fragment-based screening via MST, nsp14 constructs containing an N-terminal histidine and SUMO tags were purified. SUMO was subsequently used as a control to exclude false positives that were identified in the assay by binding to SUMO rather than nsp14. TSA and crystallisation experiments were conducted using the cleaved nsp14 construct, lacking the histag and SUMO fusion.

**Fig 1.**
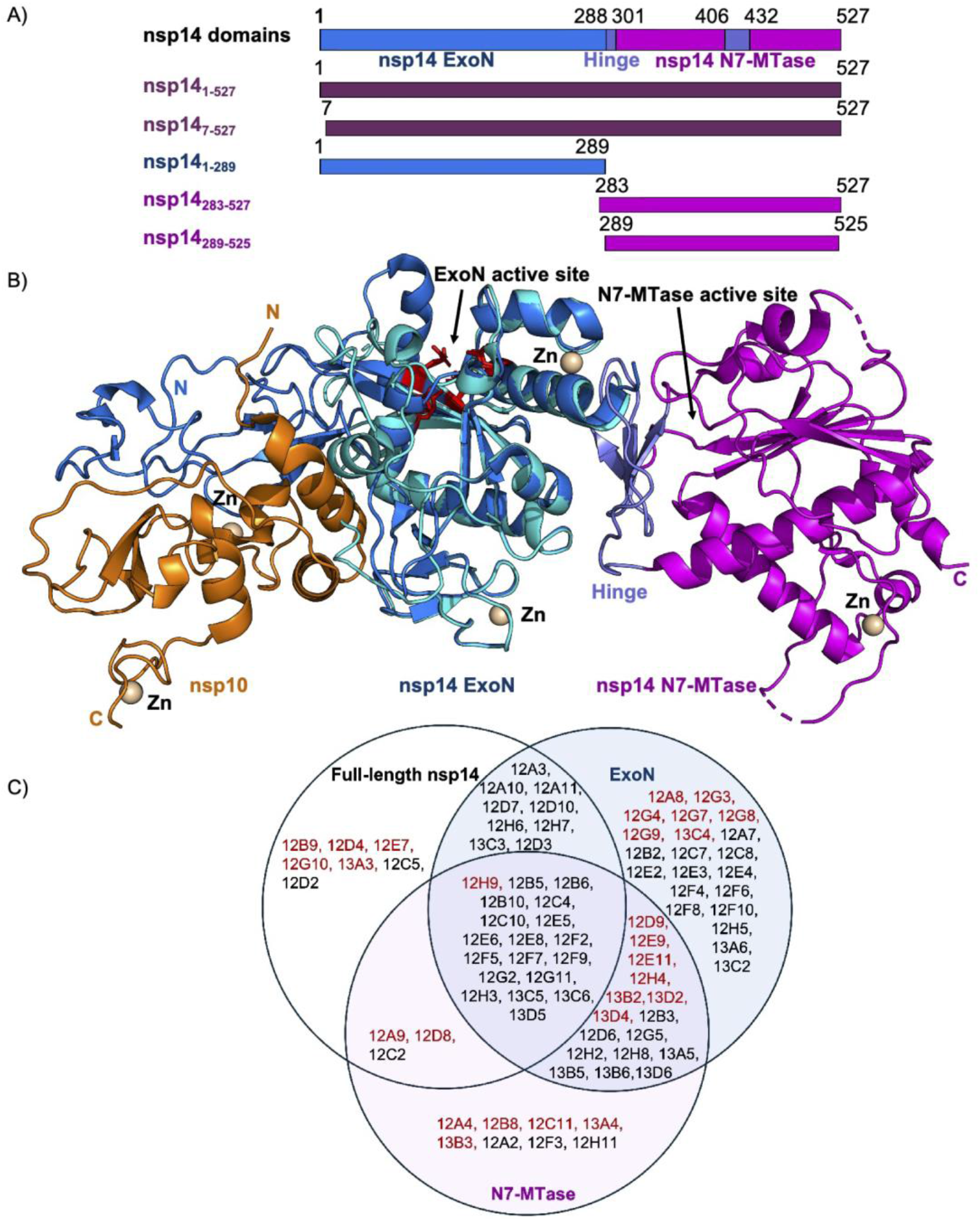
A) Bar diagram of nsp14 showing the two major ExoN (blue) and N7-MTase (magenta) domains connected via the central hinge region (purple). B) Overlay of the crystal structure of unbound nsp14 (PDB entry 7QGI) and the ExoN-nsp10 complex (PDB entry 7MC5). The three zinc finger domains are shown in light yellow, nsp10 is coloured in orange. C) Summary of MST screening results using full-length nsp14, N7-MTase and ExoN domains. The fragments listed on the chart were those identified in the binding check mode with three distinct nsp14 constructs. The fragments that were also found to bind to SUMO are shown in black and excluded from the list of positive hits. Fragment hits binding to nsp14 constructs but not to SUMO are highlighted in red.

### Identification of new nsp14 fragment hits using MST and X-ray crystallography

One hundred fragments from the Maybridge Ro3 fragment library were investigated by MST for binding to nsp14. Thirty-eight initial hits were identified for nsp14, 65 hits for the ExoN domain and 46 hits for the N7-MTase (Fig 1C). Considering that these constructs were expressed with a SUMO fusion, which may also bind fragments, we conducted a second round of MST experiments using purified SUMO. Subsequently, the hits binding to SUMO were excluded from the first-round screening results. Eight fragments were identified to bind to nsp14, 15 hits to bind to ExoN and 15 hits for N7-MTase. Some of the fragments showed binding activity for more than one nsp14 construct. For example, **12H9** bound to all three nsp14 constructs and **12A9** and **12D8** were found to bind to nsp14 and the ExoN domain, indicating that their shared binding site is localised in the ExoN domain. In total, 27 potential unique hits were identified with a success rate of 27%. All remaining fragment hits from MST were subsequently subjected to nsp14 crystal soaking and data collection at ESRF beamlines ID23-1 and ID23-2. We also recollected data for nsp14 in the absence of any fragment, which served for comparison between unbound and fragment bound electron density. This was necessary as we did not use PanDDA for automated fragment identification and did not collect diffraction data for the ground state. Out of 27 initial MST fragment hits, six resulted in nsp14-fragment complexes, a drop in the success rate to 6% (Table S1). The chemical structures of the six fragment hits are shown in Fig 2A. Data collection and refinement statistics for the nsp14-fragment complexes are summarized in Table S2.

**Fig 2.**
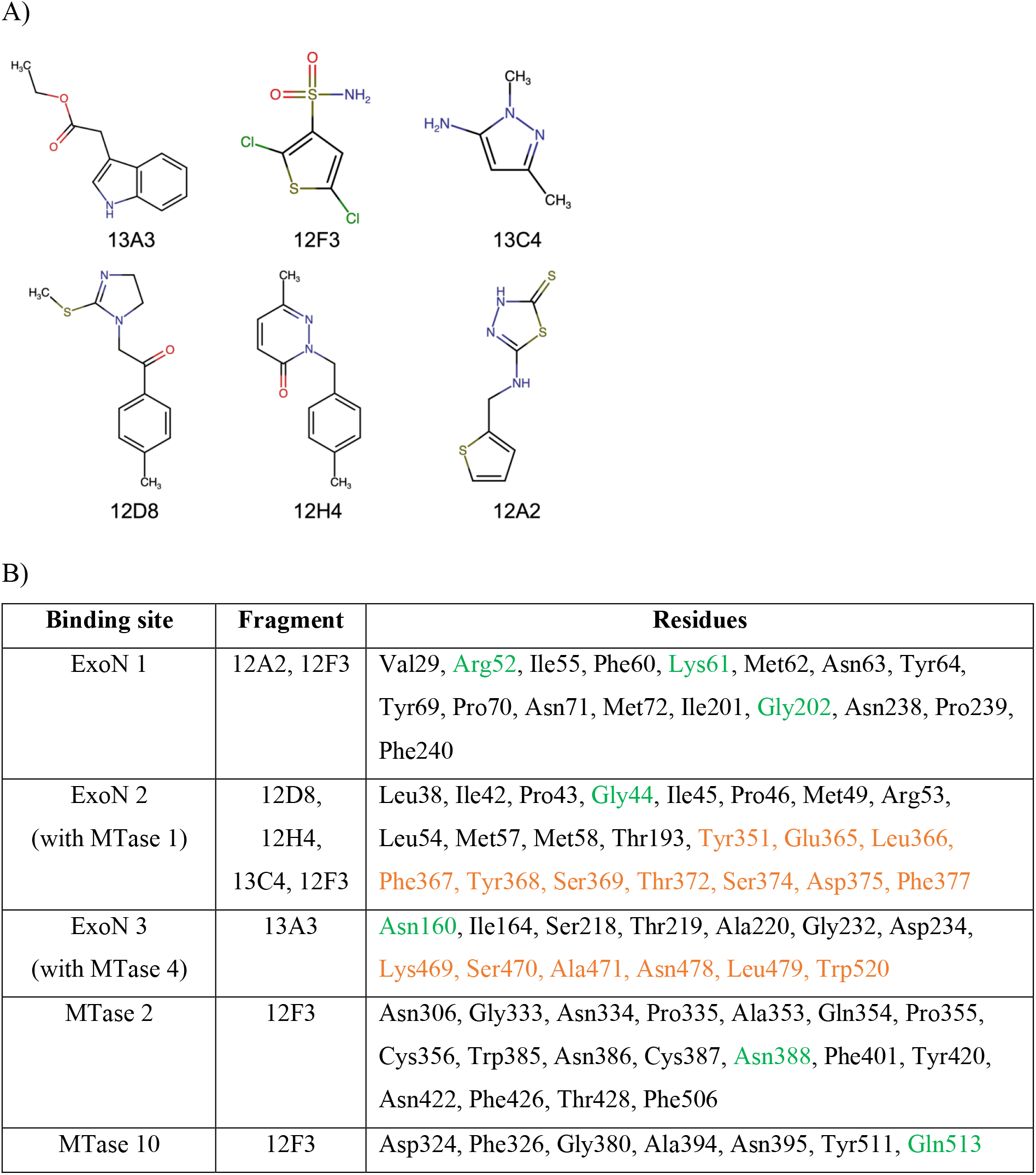
A) Chemical structures of the six fragment hits targeting SARS-CoV-2 nsp14 identified by MST and x-ray crystallography. B) Fragment binding sites on nsp14 and residues involved in the formation of these sites. Residues coloured in orange are from symmetry mates that are implicated in forming the pocket and residues shaded in green establish hydrogen bond interactions.

### Identification of fragment binding pockets on nsp14 and molecular interactions

The six chemically divergent fragment hits bind to five distinct binding sites on nsp14, with three of those being located in the ExoN domain and two being in the MTase domain (Fig 2B). We named these sites according to the location within the two domains, ExoN and MTase. Fragments **12A2** and **12F3** bind to the ExoN 1 site, whereas **12D8**, **12H4, 13C4 and 12F3** are located in the ExoN 2 site. The only fragment hit binding to the ExoN 3 site was **13A3** whereas **12F3** exclusively binds in the MTase 2 and MTase 10 site of the MTase domain. Binding of **12A2** to nsp14 induces a space group shift from *P*2_1_2_1_2_1_ to *P*2_1_ and is in the same location and conformation in both nsp14 molecules of the asymmetric unit (AU). Many fragments fall within the Ro3 with the exception of **12A2** and **12F3**, which have four instead of the preferred three hydrogen bond acceptors (Table S3).

Detailed interactions between the fragment hits and residues of nsp14 are shown in Fig. 3. Two of the fragments, **12D8 and 12H4** exclusively establish hydrophobic interactions with nsp14, whereas **12A2**, **12F3, 13C4** and **13A3** also display one or more hydrogen bonds, correlating with the larger numbers of hydrogen bond donors and acceptors being present in these fragments (Table S3). Except for the fragments **12A2** and **12F3**, which are located in the ExoN 1, MTase 2 and MTase 10 binding site, the binding pockets in the crystal structures for all other fragment hits are formed by nsp14 and one of its symmetry mates. For fragments binding to the ExoN 2 and MTase 1 sites, the number of residues from a symmetry mate involved in binding is small, consisting of only one or two residues. These sites can therefore be considered as true ExoN 2 and MTase 1 sites. Fragments in the ExoN 3 site establish three hydrophobic and one hydrogen bond interactions, as well as two hydrophobic interactions with its symmetry mate. Therefore, it is challenging to state which of the binding sites, on nsp14 or its symmetry mate, represents the true binding site, or if the combination of the two sites is required for fragment binding, therefore constituting an artificial, non-existing binding site.

**Fig 3.**
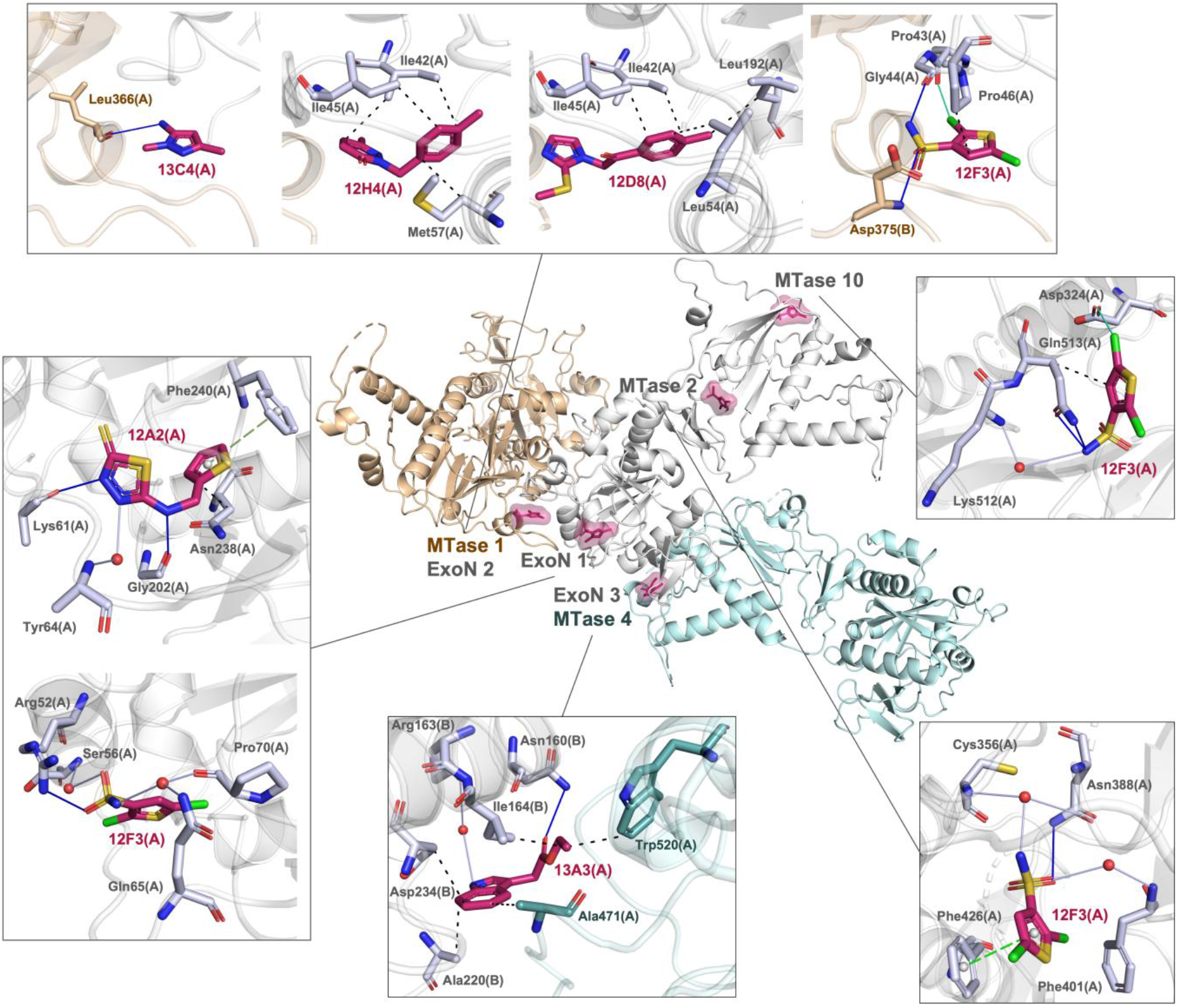
Binding sites of the fragments on nsp14 and nsp14-fragment interactions. Nsp14 is coloured in grey. The two relevant symmetry mates contributing to fragment binding are coloured in beige and cyan. The binding sites are shown in magenta surface. Hydrogen bonds are displayed as blue solid lines, hydrophobic interactions in black dashed lines, halogen bonds in green solid lines, water bridges in grey solid lines and π-stacking in green dashed lines.

### Orthogonal assays to probe the interactions between nsp14 and fragment hits

Orthogonal assays such as TSA and MST were employed to detect possible stabilisation of nsp14 or its domains and to estimate the strength of binding between nsp14 and fragment hits. We first measured TSA data for nsp14, ExoN and MTase domains in the absence and presence of DMSO. Whereas DMSO has no influence on the stability of nsp14, we observed an increase of ExoN stability in the presence of DMSO (47.4 ± 0.1°C versus 48.0 ± 0.1°C) and a slight decrease of MTase stability. However, there is no systematic trend of protein stabilisation in the presence of fragment hits for any of the nsp14 constructs tested, similar to our previous TSA experiments for other SARS-CoV-2 proteins (Table S4).

Next, MST assays were carried out for the fragment hits to estimate their K_d_ values (Table 1). Fragments **12A2** binding in the ExoN 1 site show the highest affinity with K_d_ estimates of 0.81 mM. Fragments binding to the ExoN 2 site display K_d_ estimates between 1.1 and 5.6 mM. **13A3**, binding in the ExoN 3 and MTase 4 site, has a K_d_ estimate of 3.8 mM. Interestingly, the binding pocket in the crystal is formed by nsp14 and a large number of hydrophobic contributions from a symmetry mate. This may explain the very weak binding affinity when measured in solution. In summary, K_d_ estimates are in the expected range for fragments with hits binding in the ExoN 1 site being the most promising in terms of binding affinities. Both Monolith NT.115 and Monolith X machines were used to perform MST measurements, while Monolith X also has Spectral Shift technology. The K_d_ estimates for 12H4, 12D8, and 13C4 obtained using the Monolith X machine are reported in the table, as they had a smaller error.

**Table 1.** Binding affinity of fragments and analogues to nsp14_7-527_ measured by MST.

| Fragment | Analogue | K<sub>d</sub> estimates (mM) | Labelling dye | Machine | Method |
| --- | --- | --- | --- | --- | --- |
| 12A2 |  | 0.81 ± 0.38 | RED-MALEIMIDE 2<sup>nd</sup> | Monolith NT.115 | MST |
| 12H4 |  | 1.07 [0.22-5.16] | RED-NHS 2<sup>nd</sup> | Monolith X | MST |
| 12D8 |  | 5.32 [2.84-9.96] | RED-NHS 2<sup>nd</sup> | Monolith X | Spectral Shift |
| 12F3 | 12F3 | 3.43 ± 3.40 | RED-NHS 2<sup>nd</sup> | Monolith NT.115 | MST |
|  | 6H5 | S/N=3.7 no K<sub>d</sub> | RED-NHS 2<sup>nd</sup> | Monolith NT.115 | MST |
|  | 7B2 | S/N=4.0 no K<sub>d</sub> | RED-NHS 2<sup>nd</sup> | Monolith NT.115 | MST |
|  | Q11 | 0.79 ± 0.50 | RED-NHS 2<sup>nd</sup> | Monolith NT.115 | MST |
|  | Q16 | >10 | RED-NHS 2<sup>nd</sup> | Monolith NT.115 | MST |
|  | Q17 | 2.23 ± 1.96 | RED-NHS 2<sup>nd</sup> | Monolith NT.115 | MST |
|  | 12E3 | 0.44 ± 0.37 | RED-NHS 2<sup>nd</sup> | Monolith NT.115 | MST |
| 13C4 | 13C4 | 5.63 [1.49-21.23] | RED-NHS 2<sup>nd</sup> | Monolith X | Spectral Shift |
|  | 1D10 | 1.07 ± 0.57 | RED-NHS 2<sup>nd</sup> | Monolith NT.115 | MST |
| 13A3 | 13A3 | 3.78 ± 2.69 | RED-NHS 2<sup>nd</sup> | Monolith NT.115 | MST |
|  | 3A23 | 4.23 ± 1.72 | RED-NHS 2<sup>nd</sup> | Monolith NT.115 | MST |
|  | 1G7 | no K<sub>d</sub> | RED-NHS 2<sup>nd</sup> | Monolith NT.115 | MST |

### Fragment binding sites on nsp14 in the context of functional interactions in known protein complexes

We wanted to exploit potential interference of fragment hits binding in regions of nsp14 that interact with its various binding partners. We first compared the cryo-EM nsp14-nsp10-RNA complex (PDB entry 7N0B; Liu *et al*., 2021) with our nsp14-fragment complexes overlaying their structures. Fragment hits binding in the ExoN 1 and ExoN 2 sites are located in the nsp14-nsp10 interface (Fig 4). Fragments in the ExoN 2 site are close to both the N-terminus of nsp10 and bound RNA (Fig 4). As there are significant conformational changes in the ExoN domain between unbound and nsp10 bound nsp14, fragments in ExoN 2 may stabilise the nsp10-free nsp14 conformation (Fig 4). Fragments **13A3** (ExoN 3 site) and **12F3** (MTase 1 and MTase 10 site) are not in the vicinity of either nsp10 or bound RNA. Fragment **12F3** is also in the MTase 2 SAH binding site. None of the fragments binding in the ExoN domain are close to the active site residues Asp90, Glu92, Glu191, His268 or Asp273 (DEEDh motif). In summary, two of the four binding sites are located in the nsp14-nsp10 interface and may be further developed as protein-protein interaction inhibitors.

**Fig 4.**
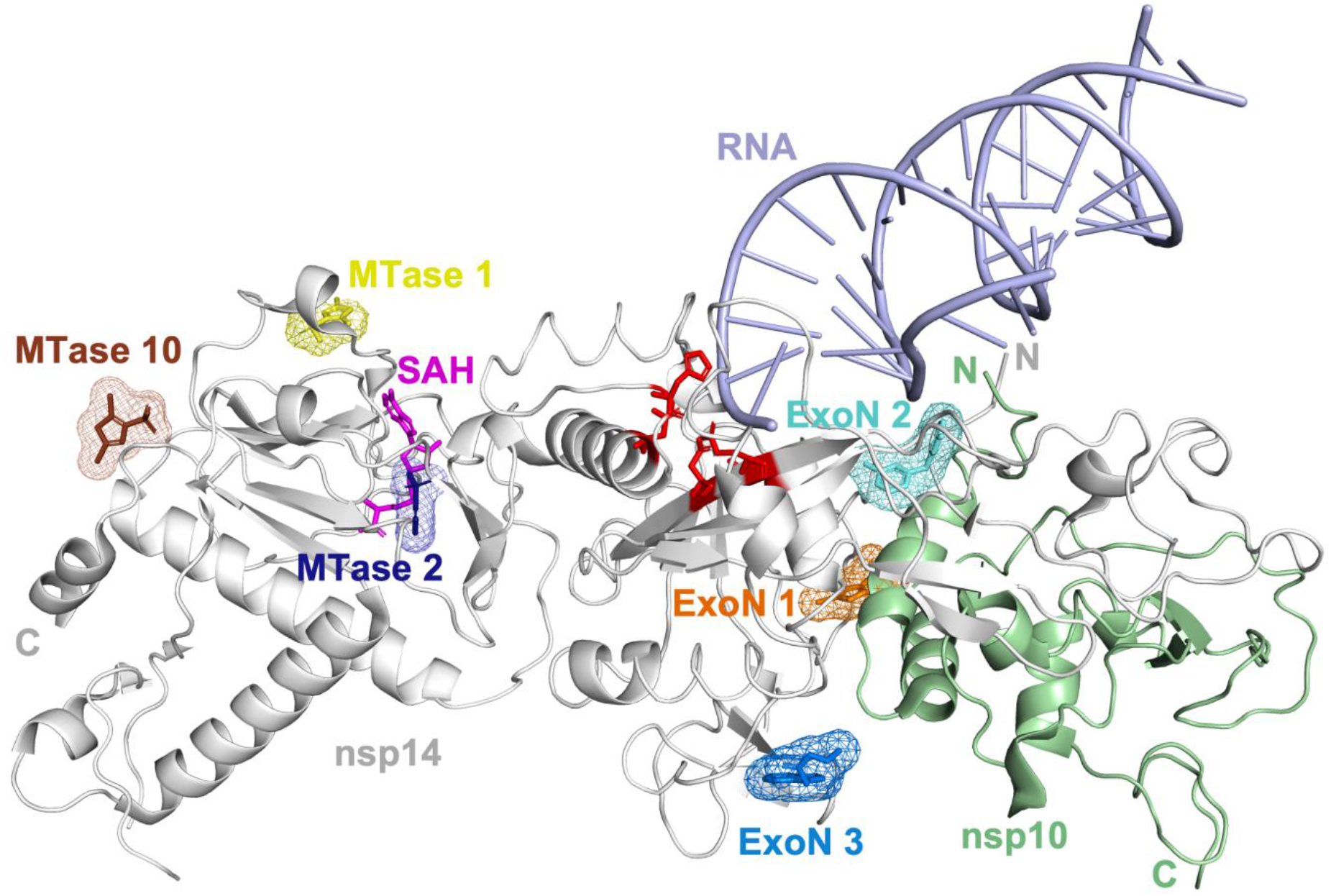
ExoN binding sites overlaid with nsp10-nsp14-RNA complex (PDB entry 7N0B). Nsp14 is shown in grey, nsp10 in green and RNA in purple. ExoN 1 site is displayed in orange, ExoN 2 in cyan, ExoN 3 in blue, MTase 1 in yellow, MTase 2 in dark blue and MTase 10 in brown. The residues in the ExoN active site are shown as sticks and coloured in red. SAH (PDB entry 7R2V) in the MTase active site is shown in magenta.

### Initial SAR-by-catalogue study

To initiate an initial SAR-by-catalogue study, we ordered commercially available fragment analogues for five of the six fragment hits. In total, we purchased 80 analogues and five additional analogues that were selected based on merging of fragments binding to the same site (Table S5). The majority of commercial analogues, 42 in total, belonged to the **13A3** scaffold. For the other hits, four to eleven analogues could be obtained, and no analogues were available for **12D8**. All analogues were subjected to crystal soaking, data collection and structure determination (summarized in Table S2). Many **13A3** analogues had a clear impact on crystal quality. After soaking, crystals with 21 distinct analogues did not diffract anymore or showed weak diffraction, even after varying analogue concentration and soaking times. For the remaining 21 analogues, only two, **3A23** and **1G7,** could be identified as analogue hits (Figure S1A). Analogues for hits **12A2**, **12H4** and those representing merged analogues of two distinct fragment hits did not bind to nsp14. For **13C4**, one out of 11 analogues did bind to nsp14 with a K_d_ of 1.07 + 0.57 mM, an approximately five-fold improvement over the original hit (Figure S1B). For fragment hit **12F3**, six out of 11 analogues did bind to nsp14, with the best analogue showing an estimated K_d_ value of 0.44 ± 0.37 mM, an almost eight-fold improvement compared to **12F3**. Rationally explaining their binding activity, even in the presence of structural data, was impossible as analogues did bind to varying pockets on nsp14 with a preference for the ExoN 2 and MTase 1 sites although other sites were also frequented such as ExoN 1, ExoN 7, MTase 2, MTase 8 and MTase 10 (Figure 7, Figure 8). In conclusion, although five-to eight-fold improvements in affinity of analogues from mM to uM could fairly easily be achieved by SAR-by-catalogue, challenges were crystal stability in the presence of certain analogue classes and “binding site hopping”, significantly complicating our SAR interpretation.

Estimates of K_d_ values were reported because some data points could be missing at high ligand concentrations in the K_d_ fit curves (Figure S4). Certain data points at high ligand concentrations (5-10 mM) had to be removed as aggregation was observed in the measurements. For binding affinity measurements with fragments that typically exhibit mM range K_d_ values, a ligand concentration greater than 10 mM would be needed to complete the K_d_ curve. However, this was challenging due to the limited solubility of compounds in the MST assay buffer. Furthermore, the DMSO concentration in measurements had to be kept at a maximum of 5% (v/v) due to its effect on protein stability. As most compound stock solutions were prepared at 200 mM in 100% DMSO, 10 mM was the highest feasible final ligand concentration in most cases.

### Identification of the most common mutations in nsp14

In order to identify important residues in nsp14 a total of 14,246,785 sequences were extracted from the GISAID database and used in this study. Using the Wuhan/WIV04/2019 sequence as reference, the nsp14 sequences were pairwise aligned to the reference, and 3435 unique types of mutations could be identified. Mutations were observed at every residue position, ranging from rare variants detected only once to highly prevalent mutations found in over eight million sequences (Fig. 6A and 6B, Fig. S2). Among the 537 residues in nsp14, 107 residues showed a very low mutation rate that exhibited a mutation count lower than 100 from more than 14 million sequences. These residues remain highly conserved, suggesting that substitutions at these positions are rarely tolerated. For example, Phe73, Arg84, Trp86, Arg310, Asp331, Gly333, Pro335, and Tyr368, which are essential for the MTase activity in SARS-CoV (Jin *et al*., 2013), also remain unchanged and highly conserved in SARS-CoV-2 (Fig. 6C). A total of 25 mutations were classified as highly common, each occurring at a frequency exceeding 0.1% (Fig. 6). Among these, the three most frequent mutations were Ile42Val, Ala394Val, and Asp222Tyr. The mutant Ile42Val was detected in nearly 60% of all SARS-CoV-2 nsp14 sequences, making it the most prevalent mutation in the dataset. Ala394Val appeared in more than 20% of sequences.

## Discussion

In this study, we used a combination of techniques to identify novel nsp14-targeting fragment hits, characterised the hits and conducted an initial SAR-by-catalogue study. At the start of this fragment screening project, structural information for SARS-CoV-2 nsp14 was not available and we turned to MST for screening. Later, X-ray and cryo-EM structures appeared in the literature for full-length nsp14, its individual domains, the nsp10-nsp14-RNA complex and several forms of the RTC (Lin *et al*., 2021; Liu *et al*., 2021; Yan *et al*., 2021; Czarna *et al*., 2022; Kottur *et al*., 2022; Moeller *et al*., 2022; Imprachim *et al*., 2023). In one study, conducting fragment-based screening via X-ray crystallography, more than 72 distinct fragment hits targeting unbound nsp14 have been described (Imprachim *et al*., 2023). In our approach, we employed a two-step cascade by first employing MST followed by X-ray crystallography. Our approach has a lower hit rate as we did not use PanDDA for fragment identification. Therefore, we could only identify fragment hits with clear positive electron density in the single crystal system, but miss low occupancy fragment hits, which are more difficult to spot in the single crystal system due to their incomplete electron density.

A comparison of our fragment hits with those previously described (Imprachim *et al*., 2023) shows that these are novel hits and that there is no overlap between these two sets. Another important question was whether we were able to identify new binding sites. By overlaying all available fragment hits, we identified a new binding site, which we named the ExoN 7 site. Due to the large number of fragment hits and associated binding sites for nsp14, we suggest introducing a naming convention for the sites, based on the location of the hits on SARS-CoV-2 nsp14, either in the ExoN domain, the central hinge region or the N7-MTase domain (Figure 5 and Table S6). Next, we explored whether, by comparing our fragments with those previously published (Imprachim *et al*., 2023), we could increase the potential for linking and merging of our new fragment hits. To investigate potential linking options, we overlaid our fragment hits binding to the ExoN domain with one representative member of previously published fragments binding in the established sites on the ExoN domain. However, the ExoN sites are located too far away from each other to be used for fragment linking.

**Fig 5.**
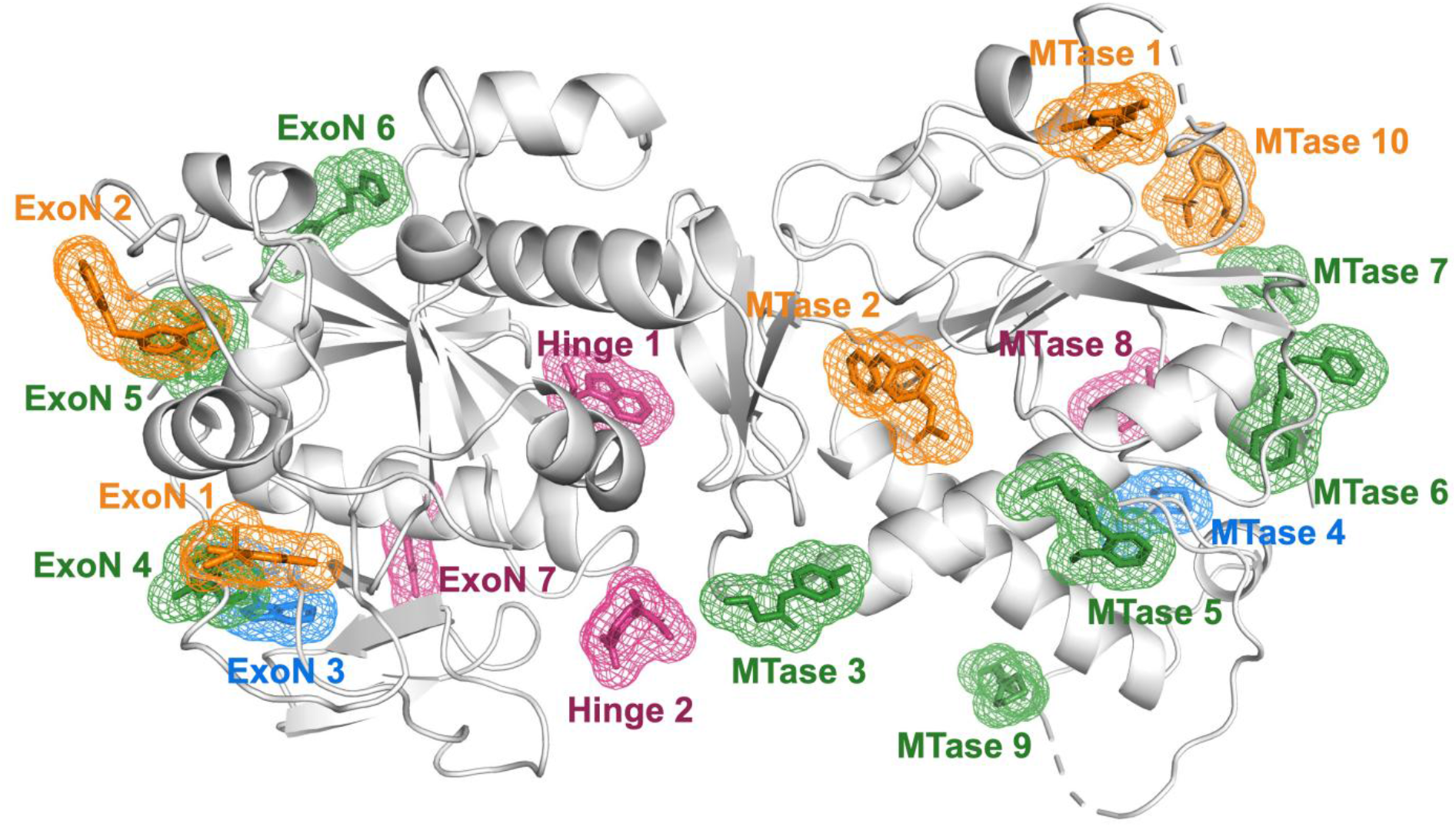
Crystal structure of nsp14 coloured in grey in the absence of its binding partner nsp10 with fragment binding sites shown as mesh. The sites that have our fragments and analogues are coloured in blue and pink, respectively. The sites that have both of our fragments and analogues are coloured in orange. The sites that only contain fragments identified by Imprachim, Yosaatmadja and Newman (2023) are shown in green. The PDB entries for the shown fragments are 5SLU, 5SLL, 5SKY, 5SL3, 5SL9, 5SM3, 5SLA, 5SM5, 5SM1, 5SL8, 5SLI, 5SLP, 9QRB, 9QRC, 9QRD, 9QXD and 9QXG.

### Ranking the existing nsp14 binding sites for their potential for further development

A total of 19 binding sites has been identified on nsp14. To assess their potential for further development into potent inhibitors, we established a range of criteria for evaluation (Table S7). One criterion was the identification of nsp14 inhibitors with proof-of-principle data, demonstrating inhibition in animal models. TDI-015051 is the only inhibitor to date that has shown such characteristics. It is a picomolar *in vitro* inhibitor against nsp14 and demonstrates antiviral efficacy in a transgenic mouse model of SARS-CoV-2 infection (Meyer *et al*., 2025). TDI-015051 binds near the SAH binding site in MTase 1, which also accommodates the cap analogue ^7^MeGpppG. Interestingly, 20 distinct fragment hits bind to this site, leading us to rank this binding site first in our evaluation.

An additional criterion was to relate the binding sites to the function of nsp14. For example, fragments binding at the nsp10-nsp14 interface may disrupt protein-protein interactions (ExoN 1, ExoN 2, ExoN 4 and ExoN 5), while those binding at the interface between nsp14 and RNA may interfere with protein-RNA interactions. Furthermore, fragment hits binding near the hinge region could potentially affect the conformational flexibility between the ExoN and MTase domains, such as at Hinge 1 and Hinge 2.

Another criterion considered the mutation frequency of residues in SARS-CoV-2 nsp14 (Figure 6). We hypothesize that residues in binding sites with high mutation frequencies will mutate under selective pressure. For example, the ExoN 6 site, which is near the ExoN active site and contains four residues with very low mutation frequencies, was ranked higher than the ExoN 2 site, which has three residues with one of the highest mutation frequencies in nsp14.

**Fig 6.**
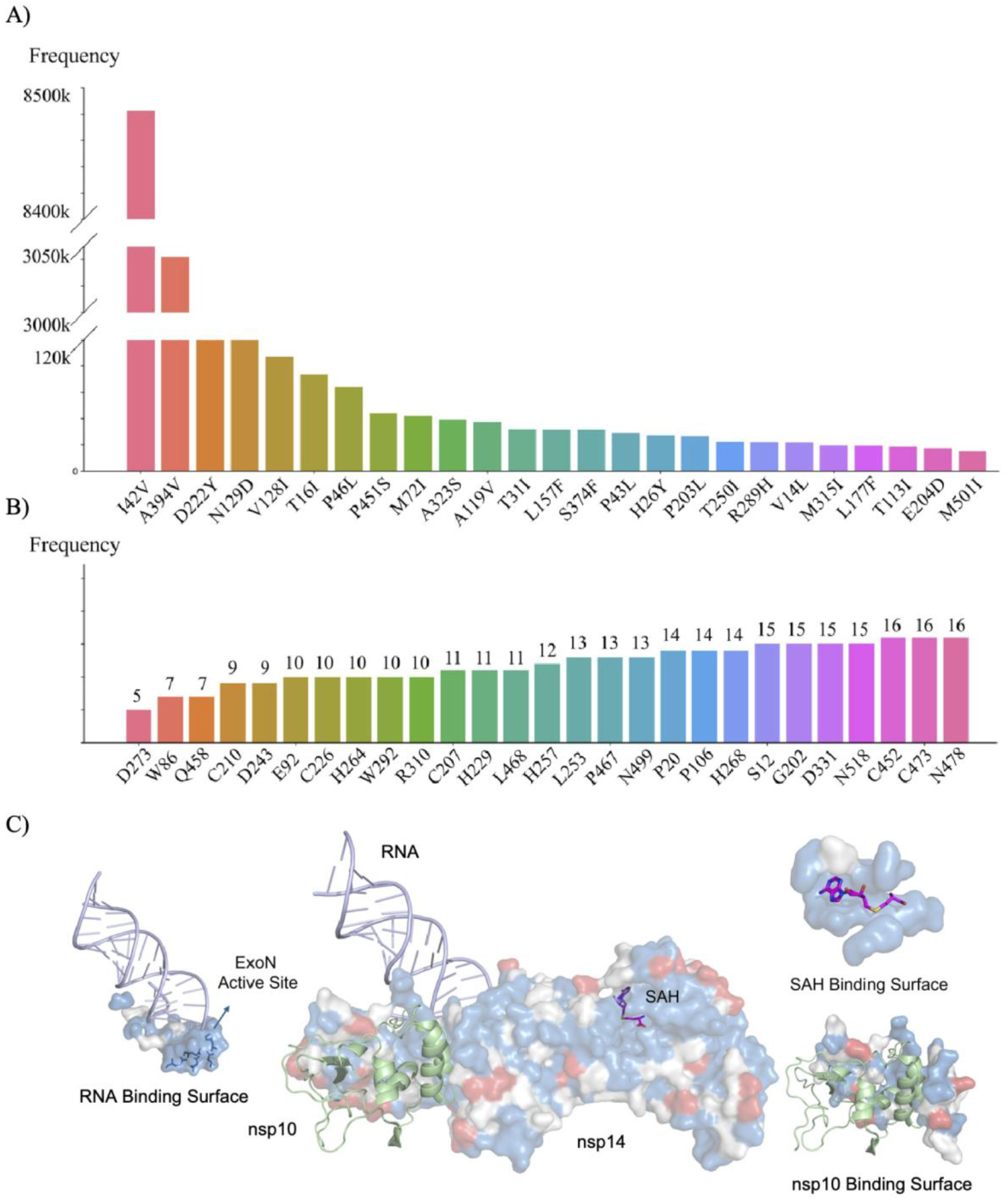
Mutations in SARS-CoV-2 nsp14. A) Mutation frequency of the top 25 mutations in SARS-CoV-2 nsp14 extracted from over 14 million sequences. B) The 25 residues with the lowest mutation frequency in nsp14. C) Surface model of nsp14 with high frequency mutations coloured in red, and low frequency amino acids shown in blue.

**Fig 7.**
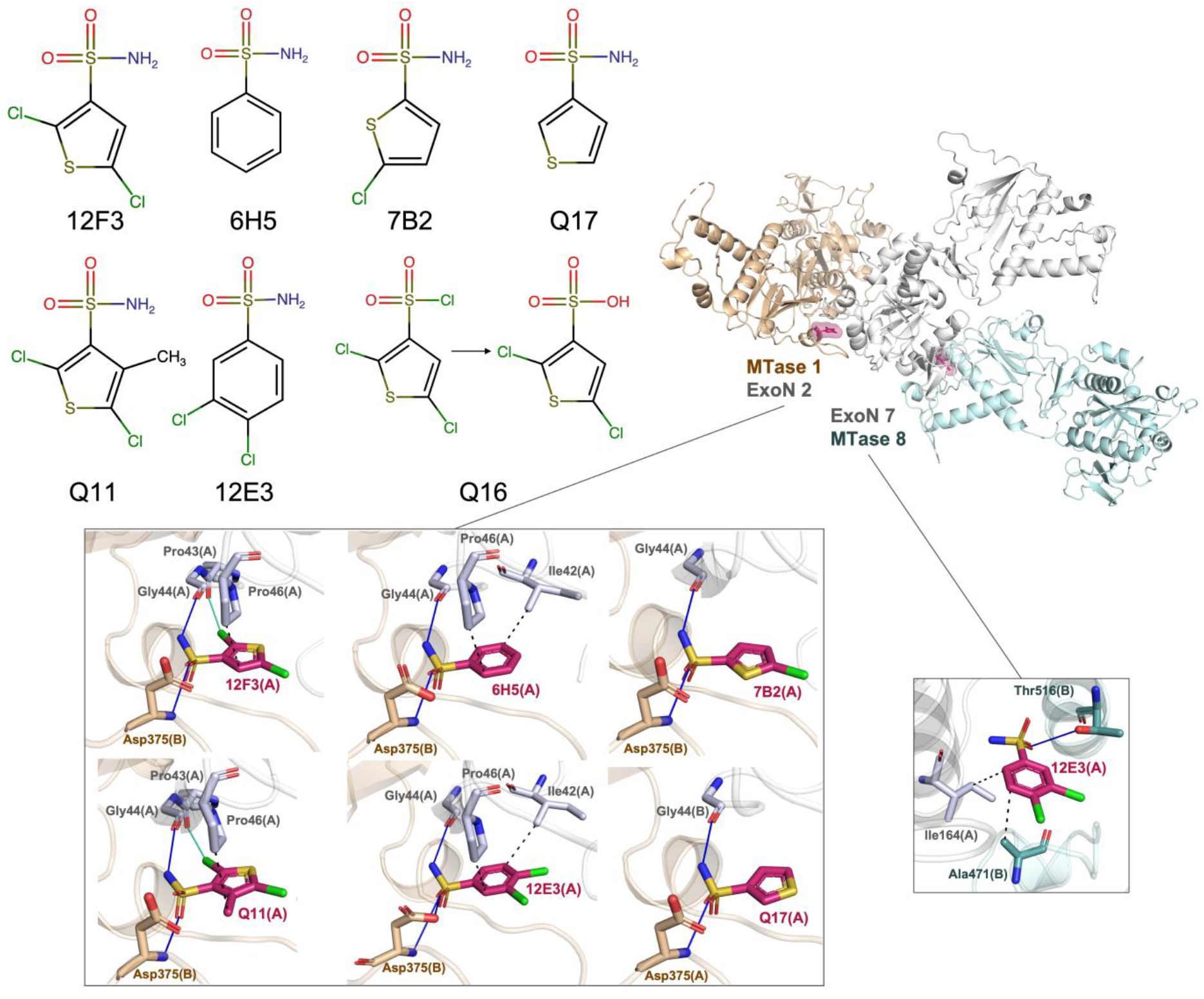
Chemical structures and interactions of 12F3 and its analogues with nsp14 residues in ExoN 2-MTase 1 and ExoN 7-MTase 8 binding sites. Nsp14 is coloured in grey. The two relevant symmetry mates contributing to fragment binding are coloured in beige and cyan. The binding sites are shown in magenta surface. Hydrogen bonds are displayed as blue solid lines, hydrophobic interactions in black dashed lines and halogen bonds in green solid lines.

**Fig 8.**
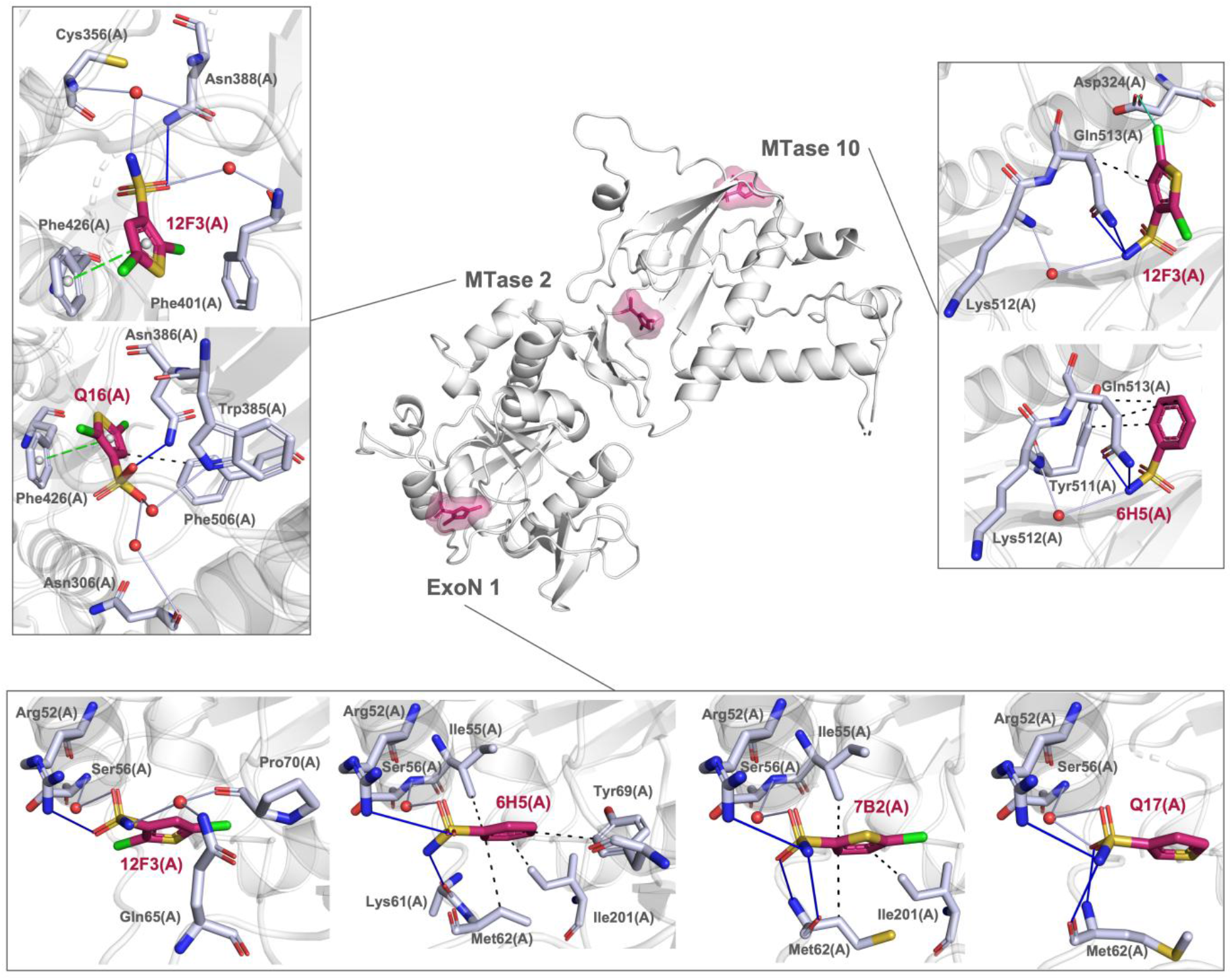
Interactions of 12F3 and its analogues with nsp14 residues in ExoN 1, MTase 2 and MTase 10 binding sites. Nsp14 is coloured in grey. The binding sites are shown in magenta surface. Hydrogen bonds are displayed as blue solid lines, hydrophobic interactions in black dashed lines, halogen bonds in green solid lines, water bridges in grey solid lines and π-stacking in green dashed lines.

A further criterion involved a protein sequence alignment between the three medically relevant CoVs SARS, SARS-CoV-2, and MERS (Fig. S3 and Table S8). Binding sites with high sequence conservation were ranked higher than those with low sequence identity. Interestingly, the three highest-ranked binding sites also showed high sequence conservation, while the lowest-ranked sites consistently exhibited low sequence identity.

Additional criteria, though considered less stringent, were also evaluated. Specifically, we assessed whether the sites represent true binding sites or if they are likely to be artifacts due to the involvement of symmetry-related molecules. Out of 19 sites, only six were formed by a combination of nsp14 and one of its symmetry mates in the crystal structure. The remaining 13 sites were determined to be “real” binding sites. We combined this criterion with the existence of orthogonal biophysical evaluations, such as MST, to support binding to a site. Estimates of K_d_ values from solution-based experiments provide further evidence of binding. Biophysical assays have been conducted for a reduced number of sites, such as ExoN 1 to ExoN 3, ExoN 7, Hinge 1, MTase 1, and MTase 2. The remaining sites require further evaluation through orthogonal assays.

Another important criterion was the number of bound fragment hits per site and the formation of subpockets. Sites that accommodate a higher number of fragments are more likely to benefit from fragment merging, and the existence of subpockets offers the potential for fragment linking. The binding sites with the highest number of fragment hits are ExoN 1, ExoN 2, Hinge 1, MTase 1, and MTase 2. Binding sites that accommodate only a limited number of fragments have reduced potential for subsequent development, as they only allow for fragment growth. Notably, the MTase 1 site, despite having 24 distinct fragments bound, was ranked low due to unfavourable factors such as high mutation frequency and low sequence conservation.

The final criterion involved the frequency of Pan Assay Interference Compounds (PAINS) per binding site. Although not directly related to the ranking of the sites, a high number of PAINS would lower the ranking of binding sites with fewer hits. However, we identified only three PAINS among all nsp14 fragments, and these occurred in sites with a high number of fragments bound. Therefore, these PAINS are not problematic when excluded from further development.

In summary, we have presented a naming and ranking system for all currently known nsp14 binding sites, which should aid in the prioritization of future development efforts for nsp14-targeting inhibitors.

### Conclusions

During our project, we identified novel fragment hits, structurally characterised their binding sites, estimated binding affinities, and contextualised our new results with existing data. By combining data from all known binding sites on nsp14, we propose a systematic naming system and ranked these binding sites based on their potential for subsequent structure-based design. Particular attention was given to mutational frequency in nsp14 and amino acid identity across the three medically relevant coronaviruses SARS, MERS, and SARS-CoV-2. It is widely accepted that viral pandemics are likely to occur again in the future, potentially originating from coronaviruses. Therefore, our work contributes to pandemic preparedness. Large conformational changes between nsp10-bound and unbound nsp14 have recently been described (Kozielski *et al*., 2025). We hypothesise that by using different approaches one may identify additional previously unidentified binding sites on nsp14.

## Data Availability

The structures of nsp14 in the presence of fragments **12A2** (PDB entry 9QS5), **12D8** (PDB entry 9QRA), **12F3** (PDB entry 9QRB), **12H4** (PDB entry 9QRC), **13A3** (PDB entry 9QRD), **13C4** (PDB entry 9QRE), **1D10** (PDB entry 9QXB), **1G7** (PDB entry 9QXC), **3A23** (PDB entry 9QXD), **6H5** (PDB entry 9QXE), **7B2** (PDB entry 9QXF), **12E3** (PDB entry 9QXG), **Q11** (PDB entry 9QXH), **Q16** (PDB entry 9QXI), and **Q17** (PDB entry 9QXK) complexes can be downloaded from the Protein Data Bank.

## Acknowledgements

This manuscript contains part of the doctoral theses of XS, QW and DD. The authors would like to thank the European Synchrotron Radiation Facility (proposals MX2488 and MX2606) and Diamond Light Source (proposal MX30393) for beamtime, and the staff of the ESRF and EMBL Grenoble for assistance and support in using beamlines ID23-1, ID23-2, ID30A-1 (MASSIF-1) and the staff of DLS beamline I03 for assistance with data collection. We thank the MRC DPFS for financial support (reference MR/X013995/1).

## Competing interests

The authors declare no competing financial interests.

## Notes

### Competing Interest Statement

The authors have declared no competing interest.

## References

Afonine, P.V., Grosse-Kunstleve, R.W., Echols, N., Headd, J.J., Moriarty, N.W., Mustyakimov, M., Terwilliger, T.C., Urzhumtsev, A., Zwart, P.H. and Adams, P.D. (2012). Towards automated crystallographic structure refinement with phenix.refine. Acta Crystallogr. D Biol. Crystallogr. 68, 352–367.

Ahmed-Belkacem, R., Hausdorff, M., Delpal, A., Sutto-Ortiz, P., Colmant, A.M.G., Touret, F., Ogando, N.S., Snijder, E.J., Canard, B., Coutard, B., Vasseur, J.J., Decroly, E. and Debart, F. (2022). Potent Inhibition of SARS-CoV-2 nsp14 N7-Methyltransferase by Sulfonamide-Based Bisubstrate Analogues. J. Med. Chem. 65, 6231–6249.

Arabi-Jeshvaghani, F., Javadi-Zarnaghi, F. and Ganjalikhany, M.R. (2023). Analysis of critical protein–protein interactions of SARS-CoV-2 capping and proofreading molecular machineries towards designing dual target inhibitory peptides. Sci. Rep. 13, 350.

Basu, S., Mak, T., Ulferts, R., Wu, M., Deegan, T., Fujisawa, R., Tan, K.W., Lim, C. T., Basier, C., Canal, B., Curran, J.F., Drury, L.S., McClure, A. W., Roberts, E.L., Weissmann, F., Zeisner, T.U., Beale, R., Cowling, V.H., Howell, M., Labib, K. and Diffley, J.F.X. (2021). Identifying SARS-CoV-2 antiviral compounds by screening for small molecule inhibitors of Nsp14 RNA cap methyltransferase. Biochem. J. 478, 2481–2497.

Baell, J.B. and Holloway, G.A. (2010). New substructure filters for removal of pan assay interference compounds (PAINS) from screening libraries and for their exclusion in bioassays. J. Med. Chem. 53, 2719–2740.

Briggs, D.C., Kandler, L., Schmidt, L., Santoni, G., and Thorn, A. (2023). Structural biology of SARS-CoV-2 accessory proteins. Crystallography Reviews. 29(1), 3–18.

Bobiļeva, O., Bobrovs, R., Kaņepe, I., Patetko, L., Kalniņš, G., Šišovs, M., Bula, A.L., Gri̅nberga, S., Borodušķis, M., Ramata-Stunda, A., Rostoks, N., Jirgensons, A., Ta̅rs, K. and Jaudzems, K. (2021). Potent SARS-CoV-2 mRNA Cap Methyltransferase Inhibitors by Bioisosteric Replacement of Methionine in SAM Cosubstrate. ACS Med. Chem. Lett. 12, 1102– 1107.

Bouvet, M., Imbert, I., Subissi, L., Gluais, L., Canard, B. and Decroly, E. (2012). RNA 3′-end mismatch excision by the severe acute respiratory syndrome coronavirus nonstructural protein nsp10/nsp14 exoribonuclease complex. Proc. Natl. Acad. Sci. U.S.A. 109, 9372–9377.

Bowler, M.W., Nurizzo, D., Barrett, R., Beteva, A., Bodin, M., Caserotto, H., Delagenière, S., Dobias, F., Flot, D., Giraud, T., Guichard, N., Guijarro, M., Lentini, M., Leonard, G.A., McSweeney, S., Oskarsson, M., Schmidt, W., Snigirev, A., Von Stetten, D., Surr, J., Svensson, O., Theveneau, P. and Mueller-Dieckmann, C. (2015). MASSIF-1: A beamline dedicated to the fully automatic characterization and data collection from crystals of biological macromolecules. J. Syn. Rad. 22, 1540–1547.

Chen, Y., Cai, H., Pan, J., Xiang, N., Tien, P., Ahola, T., and Guo, D. (2009). Functional screen reveals SARS coronavirus nonstructural protein nsp14 as a novel cap N7 methyltransferase. Proc. Natl. Acad. Sci. U.S.A. 106, 3484–3489.

Coker, J.A., Sun, R., Polzer, P.M., Romigh, T., Goins, C.M., Wang, N.S., Jung, J.U. and Stauffer, S.R. (2026) Fragment-Based Development of NSP14 Exonuclease Inhibitors Confounded by Batch-to-Batch Variability. ACS Chem. Biol. 21(3), 413–418.

Czarna, A., Plewka, J., Kresik, L., Matsuda, A., Karim, A., Robinson, C., O’Byrne, S., Cunningham, F., Georgiou, I., Wilk, P., Pachota, M., Popowicz, G., Wyatt, P. G., Dubin, G. and Pyrć, K. (2022). Refolding of lid subdomain of SARS-CoV-2 nsp14 upon nsp10 interaction releases exonuclease activity. Structure 30, 1050–1054.

Daily, J. (2016). Parasail: SIMD C library for global, semi-global, and local pairwise sequence alignments. BMC Bioinformatics 17, 81.

Emsley, P., Lohkamp, B., Scott, W.G. and Cowtan, K. (2010). Features and development of Coot. Acta Crystallogr. D Biol. Crystallogr. 66, 486–501.

Ferron, F., Subissi, L., De Morais, A.T.S., Le, N.T.T., Sevajol, M., Gluais, L., Decroly, E., Vonrhein, C., Bricogne, G., Canard, B. and Imbert, I. (2017). Structural and molecular basis of mismatch correction and ribavirin excision from coronavirus RNA. Proc. Natl. Acad. Sci. U.S.A. 115, E162–E171.

Georgiou, I., Robinson, C., O’Byrne, S.N., Matsuda, A., Grygier, P., Smith, C.D., O’Neill, S., Ahmad, S.A., Norval, S., Post, J.M., Groenewold, M., Urakova, N., Wanningen, P., Kresik, L., Plewka, J., Delpal, A., See, K., Eadsforth, T., Wierzbicka, K., Decroly, E., Saikatendu, K.S., Chang, E., Snijder, E.J., Pyrć, K., Czarna, A., Scott, D.E. and Gilbert, I.H. (2026). Crystallographic characterisation and development of bi-substrate inhibitors of coronavirus nsp14 methyltransferase. RSC Med. Chem. 17(2), 1015–1030.

Ghosh, A. and Lima, C.D. (2010). Enzymology of RNA cap synthesis. Wiley Interdisciplinary Reviews: RNA 1, 152–172.

Hampton Research. (2024). Crystal Growth 101 Literature. Available at: https://hamptonresearch.com/crystal-growth101.php.

Hsu, J.C.-C., Laurent-Rolle, M., Pawlak, J.B., Wilen, C.B. and Cresswell, P. (2021). Translational shutdown and evasion of the innate immune response by SARS-CoV-2 NSP14 protein. Proc. Natl. Acad. Sci. U.S.A. 118, p. e2101161118.

Imprachim, N., Yosaatmadja, Y. and Newman, J.A. (2023). Crystal structures and fragment screening of SARS-CoV-2 NSP14 reveal details of exoribonuclease activation and mRNA capping and provide starting points for antiviral drug development. Nucleic Acids Res. 51, 475– 487.

Jin, X., Chen, Y., Sun, Y., Zeng, C., Wang, Y., Tao, J., Wu, A., Yu, X., Zhang, Z., Tian, J. and Guo, D. (2013). Characterization of the guanine-N7 methyltransferase activity of coronavirus nsp14 on nucleotide GTP. Virus Res. 176, 45–52.

Jung, E., Soto-Acosta, R., Xie, J., Wilson, D.J., Dreis, C.D., Majima, R., Edwards, T.C., Geraghty, R.J. and Chen, L. (2022). Bisubstrate Inhibitors of Severe Acute Respiratory Syndrome Coronavirus-2 Nsp14 Methyltransferase. ACS Med. Chem. Lett. 13, 1477–1484.

Kakavandi, S., Zare, I., VaezJalali, M., Dadashi, M., Azarian, M., Akbari, A., Ramezani Farani, M., Zalpoor, H. and Hajikhani, B. (2023). Structural and non-structural proteins in SARS-CoV-2: potential aspects to COVID-19 treatment or prevention of progression of related disease. Cell Commun. Signal. 21, 110.

Kasprzyk, R., Spiewla, T.J., Smietanski, M., Golojuch, S., Vangeel, L., De Jonghe, S., Jochmans, D., Neyts, J., Kowalska, J. and Jemielity, J. (2021). Identification and evaluation of potential SARS-CoV-2 antiviral agents targeting mRNA cap guanine N7-Methyltransferase. Antiviral Res. 193, 105142.

Khare, S., Gurry, C., Freitas, L., Schultz, M.B., Bach, G., Diallo, A., Akite, N., Ho, J., Lee, R.T.C., Yeo, W., GISAID Core Curation Team and Maurer-Stroh, S. (2021). Perspectives: GISAID’s Role in Pandemic Response. China CDC Weekly 3(49), 1049–1051.

Kottur, J., Rechkoblit, O., Quintana-Feliciano, R., Sciaky, D. and Aggarwal, A.K. (2022). High-resolution structures of the SARS-CoV-2 N7-methyltransferase inform therapeutic development. Nat. Struct. Mol. Biol. 29, 850–853.

Kozielski, F., Sele, C., Talibov, V.O., Lou, J., Dong, D., Wang, Q., Shi, X., Nyblom, M., Rogstam, A., Krojer, T., Fisher, Z. and Knecht, W. (2022). Identification of fragments binding to SARS-CoV-2 nsp10 reveals ligand-binding sites in conserved interfaces between nsp10 and nsp14/nsp16. RSC Chem. Biol. 3, 44–55.

Kozielski, F., Fisher, S.Z., Ma, S., Busaidi, F.A., Krupinska, E., Nyblom, M., Sele, C., Sullivan, H.M., Krojer, T. and Knecht, W. (2025). Structural basis for small molecule binding to the SARS-CoV-2 nsp10–nsp14 ExoN complex. Nucleic Acids Research. 53(14).

Lagorce, D., Sperandio, O., Baell, J.B., Miteva, M.A. and Villoutreix, B.O. (2015). FAF-Drugs3: A web server for compound property calculation and chemical library design. Nucleic Acids Res. 43, W200–W207.

Liebschner, D., Afonine, P.V., Baker, M.L., Bunkóczi, G., Chen, V.B., Croll, T.I., Hintze, B., Hung, L.-W., Jain, S., McCoy, A.J., Moriarty, N.W., Oeffner, R.D., Poon, B.K., Prisant, M. G., Read, R.J., Richardson, J.S., Richardson, D.C., Sammito, M.D., Sobolev, O.V., Stockwell, D.H., Terwilliger, T.C., Urzhumtsev, A.G., Videau, L.L., Williams, C.J. and Adams, P.D. (2019). Macromolecular structure determination using X-rays, neutrons and electrons: recent developments in Phenix. Acta Crystallogr. D Biol. Crystallogr. 75, 861–877.

Lin, S., Chen, H., Chen, Z., Yang, F., Ye, F., Zheng, Y., Yang, J., Lin, X., Sun, H., Wang, L., Wen, A., Dong, H., Xiao, Q., Deng, D., Cao, Y. and Lu, G. (2021). Crystal structure of SARS-CoV-2 nsp10 bound to nsp14-ExoN domain reveals an exoribonuclease with both structural and functional integrity. Nucleic Acids Res. 49, 5382–5392.

Liu, C., Shi, W., Becker, S.T., Schatz, D.G., Liu, B. and Yang, Y. (2021). Structural basis of mismatch recognition by a SARS-CoV-2 proofreading enzyme. Science 373, 1142–1146.

Mariano, G., Farthing, R.J., Lale-Farjat, S.L.M., and Bergeron, J.R.C. (2020). Structural Characterization of SARS-CoV-2: Where We Are, and Where We Need to Be. Front. Mol. Biosci. 7, 2020.

McCoy, A.J., Grosse-Kunstleve, R.W., Adams, P.D., Winn, M.D., Storoni, L.C. and Read, R.J. (2007). Phaser crystallographic software. J. Appl. Crystallogr. 40, 658–674.

Meyer, C., Garzia, A., Miller, M. W., Huggins, D. J., Myers, R. W., Hoffmann, H. H., Ashbrook, A. W., Jannath, S. Y., Liverton, N., Kargman, S., Zimmerman, M., Nelson, A. M., Sharma, V., Dolgov, E., Cangialosi, J., Penalva-Lopez, S., Alvarez, N., Chang, C. W., Oswal, N., Gonzalez, I.,…Tuschl, T. (2025). Small-molecule inhibition of SARS-CoV-2 NSP14 RNA cap methyltransferase. Nature 637(8048), 1178–1185.

Minskaia, E., Hertzig, T., Gorbalenya, A.E., Campanacci, V., Cambillau, C., Canard, B., and Ziebuhr, J. (2006). Discovery of an RNA virus 3′-5′ exoribonuclease that is critically involved in coronavirus RNA synthesis. Proc. Natl. Acad. Sci. U.S.A. 103, 5108–5113.

Moeller, N.H., Shi, K., Demir, Ö., Belica, C., Banerjee, S., Yin, L., Durfee, C., Amaro, R.E. and Aihara, H. (2022). Structure and dynamics of SARS-CoV-2 proofreading exoribonuclease ExoN. Proc. Natl. Acad. Sci. U.S.A. 119, e2106379119.

Monaco, S., Gordon, E., Bowler, M.W., Delagenière, S., Guijarro, M., Spruce, D., Svensson, O., McSweeney, S.M., McCarthy, A.A., Leonard, G. and Nanao, M.H. (2013). Automatic processing of macromolecular crystallography X-ray diffraction data at the ESRF. J. Appl. Crystallogr. 46, 804–810.

Ogando, N. S., Zevenhoven-Dobbe, J. C., van der Meer, Y., Bredenbeek, P. J., Posthuma, C. C. and Snijder, E. J. (2020). The Enzymatic Activity of the nsp14 Exoribonuclease Is Critical for Replication of MERS-CoV and SARS-CoV-2. Journal of Virology 94(23), e01246–20.

Pearson, L.-A., Green, C.J., Lin, D., Petit, A.-P., Gray, D.W., Cowling, V.H. and Fordyce, E.A.F. (2021). Development of a High-Throughput Screening Assay to Identify Inhibitors of the SARS-CoV-2 Guanine-N7-Methyltransferase Using RapidFire Mass Spectrometry. SLAS Discov. 26, 749–756.

Rona, G., Zeke, A., Miwatani-Minter, B., de Vries, M., Kaur, R., Schinlever, A., Garcia, S.F., Goldberg, H.V., Wang, H., Hinds, T.R., Bailly, F., Zheng, N., Cotelle, P., Desmaële, D., Landau, N.R., Dittmann, M. and Pagano, M. (2022). The NSP14/NSP10 RNA repair complex as a Pan-coronavirus therapeutic target. Cell Death Differ. 29, 285–292.

Schake, P., Bolz, S.N., Linnemann, K. and Schroeder, M. (2025). PLIP 2025: introducing protein-protein interactions to the protein-ligand interaction profiler. Nucleic Acids Research 53, 463–465.

Subissi, L., Imbert, I., Ferron, F., Collet, A., Coutard, B., Decroly, E. and Canard, B. (2014). SARS-CoV ORF1b-encoded nonstructural proteins 12-16: Replicative enzymes as antiviral targets. Antiviral Research. 101, 122–130.

Svensson, O., Malbet-Monaco, S., Popov, A., Nurizzo, D. and Bowler, M.W. (2015). Fully automatic characterization and data collection from crystals of biological macromolecules. Acta Crystallogr. D Biol. Crystallogr. 71, 1757–1767.

Vonrhein, C., Flensburg, C., Keller, P., Sharff, A., Smart, O., Paciorek, W., Womack, T. and Bricogne, G. (2011). Data processing and analysis with the autoPROC toolbox. Acta Crystallogr. D Biol. Crystallogr. 67, 293–302.

Winn, M.D., Ballard, C.C., Cowtan, K.D., Dodson, E.J., Emsley, P., Evans, P.R., Keegan, R.M., Krissinel, E.B., Leslie, A.G. W., McCoy, A., McNicholas, S.J., Murshudov, G.N., Pannu, N.S., Potterton, E.A., Powell, H.R., Read, R.J., Vagin, A. and Wilson, K.S. (2011). Overview of the CCP4 suite and current developments. Acta Crystallogr. D Biol. Crystallogr. 67, 235– 242.

World Health Organization (2026). Number of COVID-19 cases reported to WHO (cumulative total). Available at: https://data.who.int/dashboards/covid19/cases?n=c.

Winter, G. (2010). xia2: an expert system for macromolecular crystallography data reduction. J. Appl. Crystallogr. 43, 186–190.

Winter, G., Waterman, D.G., Parkhurst, J.M., Brewster, A.S., Gildea, R.J., Gerstel, M., Fuentes-Montero, L., Vollmar, M., Michels-Clark, T., Young, I.D., Sauter, N.K. and Evans, G. (2018). DIALS: implementation and evaluation of a new integration package. Acta Crystallogr. D Biol. Crystallogr. 74, 85–97.

Yan, L., Yang, Y., Li, M., Zhang, Y., Zheng, L., Ge, J., Huang, Y.C., Liu, Z., Wang, T., Gao, S., Zhang, R., Huang, Y.Y., Guddat, L.W., Gao, Y., Rao, Z. and Lou, Z. (2021). Coupling of N7-methyltransferase and 3′-5′ exoribonuclease with SARS-CoV-2 polymerase reveals mechanisms for capping and proofreading. Cell 184, 3474–3485.

