## Supplementary Material for "Mapping the Fragment-Binding Landscape of SARS-CoV-2 nsp14 for Antiviral Drug Discovery"

**Table S1.** Summary of screening cascade for fragments binding to nsp14 using a combination of MST and X-ray crystallography as primary and secondary screen, respectively.

|  | **Full-length nsp14**<br>**(No. hits)** | **ExoN**<br>**(No. hits)** | **N7-MTase**<br>**(No. hits)** |
| --- | --- | --- | --- |
| **1<sup>st</sup> round MST** | 38 | 65 | 46 |
| **2<sup>nd</sup> round MST (SUMO)** | 8 | 15 | 15 |
| **Unique hits** | 5 | 5 | 5 |
| **X-ray crystallography** | 6 | N/A | N/A |

**Table S2.** Data collection, structure determination and refinement statistics for nsp14-fragment and analogue complexes. Data in parenthesis correspond to the highest resolution shell.

| Parameters | Nsp14-12D8 | Nsp14-12H4 | Nsp14-13A3 |
| --- | --- | --- | --- |
| Wavelength [Å] | 0.87313 |  | 0.88560 |
| Resolution range [Å] | 69.334 - 2.294<br>(2.334 - 2.294) | 61.101 - 2.404<br>(2.49 - 2.404) | 48.46 - 1.91<br>(1.95 - 1.91) |
| Space group | $P2_12_12_1$ | | |
| Unit cell parameters [Å, °] | $a=68.115$,<br>$b=68.392$,<br>$c=138.667$,<br>$\alpha=\beta=\gamma=90$ | $a=67.992$,<br>$b=68.431$,<br>$c=139.284$,<br>$\alpha=\beta=\gamma=90$ | $a=67.489$,<br>$b=68.031$,<br>$c=138.117$,<br>$\alpha=\beta=\gamma=90$ |
| Molecules per asymmetric unit | 1 |  |  |
| Total reflections | 151628 (7740) | 171896<br>(18106) | 212963<br>(12700) |
| Unique reflections | 28658 (1428) | 25907 (2517) | 48906 (3260) |
| Multiplicity | 5.3 (5.4) | 6.6 (6.8) | 4.4 (3.9) |
| Completeness [%] | 95.6 (98.4) | 99.5 (98.3) | 97.2 (97.1) |
| Mean I/sigma(I) | 7.0 (1.5) | 10.3 (1.3) | 9.6 (1.6) |
| Wilson B-factor (Å<sup>2</sup>) | 30.3 | 49.3 | 31.4 |
| R<sub>meas</sub> [%] | 22.9 (204.3) | 16.5 (164.2) | 9.2 (84.2) |
| CC<sub>1/2</sub> | 0.967 (0.698) | 0.996 (0.520) | 0.996 (0.628) |
| Reflections used in refinement | 28445 (2880) | 25897 (2515) | 48732 (4791) |
| R<sub>cryst</sub> / R<sub>free</sub> [%] | 19.1 (25.0) /<br>25.0 (29.9) | 19.4 (33.9) /<br>24.0 (39.9) | 19.1 (28.0) /<br>22.8 (31.2) |
| Total no. of non-hydrogen atoms (protein) | 3702 | 3606 | 3850 |
| No. of protein/ligand/solvent residues | 446 / 9 / 152 | 445 / 9 / 64 | 442 / 9 / 335 |
| RMSD bond length, bond angles [Å, °] | 0.012 / 1.43 | 0.012 / 1.40 | 0.012 / 1.13 |
| Ramachandran favoured/allowed/outliers/rotamer outliers [%] | 96.1 / 3.4 / 0.5<br>/ 2.1 | 95.4 / 4.1 / 0.5<br>/ 3.0 | 97.0 / 2.3 / 0.7<br>/ 0.83 |
| Clashscore | 3.6 | 3.2 | 3.7 |
| Average B-factor/protein/ligands/solvent | 35.7 / 35.5 /<br>50.5 / 36.8 | 57.2 / 57.0 /<br>72.4 / 57.7 | 38.9 / 38.0 /<br>57.6 / 45.6 |

**Continuation Table S2.** Data collection, structure determination and refinement statistics for nsp14-fragment and analogue complexes. Data in parenthesis correspond to the highest resolution shell.

| Parameters | Nsp14-13C4 | Nsp14-12F3 | Nsp14-12A2 |
| --- | --- | --- | --- |
| Wavelength [Å] | 0.87313 |  | 0.88560 |
| Resolution range [Å] | 69.477 - 2.298<br>(2.380 - 2.298) | 61.092 - 1.935<br>(2.005 - 1.935) | 49.68 – 2.20<br>(2.28 – 2.20) |
| Space group | P2<sub>1</sub>2<sub>1</sub>2<sub>1</sub> |  | P2<sub>1</sub> |
| Unit cell parameters [Å, °] | *a*=68.337,<br>*b*=68.751,<br>*c*=138.953,<br>$\alpha=\beta=\gamma=90$ | *a*=67.982,<br>*b*=68.38,<br>*c*=139.268,<br>$\alpha=\beta=\gamma=90$ | *a*=63.773,<br>*b*=138.145,<br>*c*=71.515,<br>$\alpha=90, \beta=90.005, \gamma=90$ |
| Molecules per asymmetric unit | 1 |  | 2 |
| Total reflections | 315211 (15742) | 333503 (22014) | 118997 (9461) |
| Unique reflections | 28859 (1454) | 49211 (3243) | 54220 (4647) |
| Multiplicity | 10.9 (10.8) | 6.8 (6.8) | 2.2 (2.0) |
| Completeness [%] | 96.5 (100.0) | 99.9 (100.0) | 87.0 (90.5) |
| Mean I/sigma(I) | 7.8 (1.7) | 13.2 (1.5) | 8.3 (1.5) |
| Wilson B-factor (Å<sup>2</sup>) | 31.8 | 33.3 | 37.6 |
| R<sub>meas</sub> [%] | 40.9 (511.3) | 10.5 (167.8) | 8.9 (55.6) |
| CC<sub>1/2</sub> | 0.989 (0.467) | 0.998 (0.673) | 0.987 (0.800) |
| Reflections used in refinement | 28580 (2795) | 48298 (4603) | 54114 (5596) |
| R<sub>cryst</sub> / R<sub>free</sub> [%] | 19.5 (27.7) /<br>25.0 (30.9) | 19.3 (29.0) / 23.3<br>(30.7) | 20.0 (32.9) /<br>24.9 (35.8) |
| Total no. of non-hydrogen atoms (protein) | 3705 | 3952 | 7138 |
| No. of protein/ligand/solvent residues | 452 / 12 / 116 | 453 / 11 / 284 | 891 / 25 / 127 |
| RMSD bonds length, bond angles [Å, °] | 0.012 / 1.20 | 0.012 / 1.08 | 0.012 / 1.26 |
| Ramachandran favoured/allowed/outliers/rotamer outliers [%] | 95.5 / 4.3 / 0.2 /<br>2.5 | 97.8 / 2.3 / 0.0 /<br>0.3 | 94.5 / 4.7 / 0.8<br>/ 1.6 |
| Clashscore | 3.3 | 3.0 | 6.2 |
| Average B-factor/protein/ligands/solvent | 39.1 / 38.6 /<br>75.3 / 39.4 | 38.8 / 38.0 / 60.8<br>/ 43.3 | 44.2 / 43.8 /<br>72.6 / 44.6 |

**Continuation Table S2.** Data collection, structure determination and refinement statistics for nsp14-fragment and analogue complexes. Data in parenthesis correspond to the highest resolution shell.

| Parameters | Nsp14-6H5 | Nsp14-7B2 | Nsp14-12E3 |
| --- | --- | --- | --- |
| Wavelength [Å] | 0.9763 |  |  |
| Resolution range [Å] | 45.23 - 1.93<br>(2.00 - 1.93) | 48.63 - 2.25<br>(2.33 - 2.25) | 60.95 - 2.3<br>(2.38 - 2.3) |
| Space group | $P2_12_12_1$ | | |
| Unit cell parameters [Å, °] | a=67.361,<br>b=68.041,<br>c=138.152,<br>$\alpha=\beta=\gamma=90$ | a=67.400,<br>b=68.148,<br>c=138.828,<br>$\alpha=\beta=\gamma=90$ | a=67.163,<br>b=67.812,<br>c=139.082,<br>$\alpha=\beta=\gamma=90$ |
| Molecules per asymmetric unit | 1 |  |  |
| Total reflections | 655652<br>(44303) | 422178<br>(39924) | 392610<br>(39912) |
| Unique reflections | 48589 (3228) | 31154 (2825) | 29033 (2824) |
| Multiplicity | 13.5 (13.7) | 13.6 (14.1) | 13.5 (14.1) |
| Completeness [%] | 100 (100) | 100 (100) | 100 (99.9) |
| Mean I/sigma(I) | 14.4 (1.4) | 13.7 (1.4) | 11.4 (1.5) |
| Wilson B-factor (Å<sup>2</sup>) | 30.0 | 38.8 | 42.2 |
| R<sub>meas</sub> [%] | 14.1 (182.6) | 16.0 (187.6) | 20.1 (204.1) |
| CC<sub>1/2</sub> | 0.999 (0.838) | 0.998 (0.91) | 0.998 (0.802) |
| Reflections used in refinement | 48133 (4777) | 30887 (2889) | 28931 (2859) |
| R<sub>cryst</sub> / R<sub>free</sub> [%] | 19.9 (28.7) /<br>23.2 (33.3) | 20.4 (41.4) /<br>25.0 (45.7) | 20.8 (32.0) /<br>25.7 (39.3) |
| Total no. of non-hydrogen atoms (protein) | 4023 | 3833 | 3723 |
| No. of protein/ligand/solvent residues | 453 / 14 / 331 | 454 / 12 / 181 | 452 / 13 / 116 |
| RMSD bond length, bond angles [Å, °] | 0.012 / 1.17 | 0.012 / 1.31 | 0.012 / 1.35 |
| Ramachandran favoured/allowed/outliers/rotamer outliers [%] | 97.08 / 2.47 /<br>0.45 / 1.31 | 95.29 / 3.59 /<br>1.12 / 2.93 | 93.02 / 5.86 /<br>1.13 / 3.54 |
| Clashscore | 3.67 | 2.83 | 5.48 |
| Average B-factor/protein/ligands/solvent | 37.99 / 36.94 /<br>65.31 / 41.91 | 46.41 / 45.91 /<br>71.85 / 46.97 | 51.27 / 50.95 /<br>70.65 / 50.23 |

**Continuation Table S2.** Data collection, structure determination and refinement statistics for nsp14-fragment and analogue complexes. Data in parenthesis correspond to the highest resolution shell.

| Parameters | Nsp14-Q11 | Nsp14-Q16 | Nsp14-Q17 |
| --- | --- | --- | --- |
| Wavelength [Å] | 0.9763 |  |  |
| Resolution range [Å] | 60.92 - 2.1<br>(2.16 - 2.1) | 38.99 - 2.14<br>(2.2 - 2.14) | 61.05 – 1.91<br>(1.95 – 1.91) |
| Space group | $P2_12_12_1$ | | |
| Unit cell parameters [Å, °] | $a=67.759$,<br>$b=67.885$,<br>$c=138.030$,<br>$\alpha=\beta=\gamma=90$ | $a=66.623$,<br>$b=66.756$,<br>$c=138.661$,<br>$\alpha=\beta=\gamma=90$ | $a=67.315$,<br>$b=68.062$,<br>$c=138.122$,<br>$\alpha=\beta=\gamma=90$ |
| Molecules per asymmetric unit | 1 |  |  |
| Total reflections | 510771<br>(42277) | 469168<br>(37933) | 681269<br>(46375) |
| Unique reflections | 38001 (3075) | 34905 (2803) | 50085 (3336) |
| Multiplicity | 13.4 (13.7) | 13.4 (13.5) | 13.6 (13.9) |
| Completeness [%] | 100 (100) | 99.9 (99.9) | 100 (100) |
| Mean I/sigma(I) | 10.1 (1.5) | 15.1 (1.5) | 14.8 (1.6) |
| Wilson B-factor (Å<sup>2</sup>) | 31.6 | 38.3 | 28.7 |
| R<sub>meas</sub> [%] | 23.2 (203.7) | 13.1 (177.2) | 14.9 (183.1) |
| CC<sub>1/2</sub> | 0.997 (0.814) | 0.999 (0.916) | 0.999 (0.795) |
| Reflections used in refinement | 37619 (3728) | 34281 (3389) | 49943 (4929) |
| R<sub>cryst</sub> / R<sub>free</sub> [%] | 21.4 (30.7) /<br>25.7 (40.0) | 22.8 (33.0) /<br>24.7 (30.5) | 19.5 (31.8) /<br>23.3 (39.2) |
| Total no. of non-hydrogen atoms<br>(protein) | 3843 | 3651 | 4016 |
| No. of protein/ligand/solvent residues | 452 / 11 / 181 | 447 / 8 / 125 | 454 / 9 / 345 |
| RMSD bond length, bond angles<br>[Å, °] | 0.012 / 1.16 | 0.011 / 1.07 | 0.012 / 1.15 |
| Ramachandran<br>favoured/allowed/outliers/rotamer<br>outliers [%] | 95.50 / 3.38 /<br>1.13 / 1.84 | 94.97 / 5.03 /<br>0.00 / 1.94 | 97.76 / 2.02 /<br>0.22 / 1.05 |
| Clashscore | 5.51 | 4.43 | 2.82 |
| Average B-factor/protein/ligands/solvent | 41.72 / 41.18 /<br>67.70 / 42.58 | 56.65 / 56.45 /<br>75.29 / 57.41 | 36.42 / 35.55 /<br>62.09 / 41.17 |

**Continuation Table S2.** Data collection, structure determination and refinement statistics for nsp14-fragment and analogue complexes. Data in parenthesis correspond to the highest resolution shell.

| Parameters | Nsp14-3A23 | Nsp14-1G7 | Nsp14-1D10 |
| --- | --- | --- | --- |
| Wavelength [Å] | 0.965460 | 0.9763 |  |
| Resolution range [Å] | 48.42 - 2.00<br>2.07 - 2.00) | 39.36 – 1.98<br>(2.05 – 1.98) | 48.44 - 1.84<br>(1.91 - 1.84) |
| Space group | $P2_12_12_1$ | | |
| Unit cell parameters [Å, °] | $a=67.541,$<br>$b=67.838,$<br>$c=138.269,$<br>$\alpha=\beta=\gamma=90$ | $a=67.66,$<br>$b=67.89,$<br>$c=138.03,$<br>$\alpha=\beta=\gamma=90$ | $a=67.09,$<br>$b=67.77,$<br>$c=138.55,$<br>$\alpha=\beta=\gamma=90$ |
| Molecules per asymmetric unit | 1 |  |  |
| Total reflections | 157652 (11386) | 608917 (30328) | 759556 (38863) |
| Unique reflections | 42050 (3137) | 45133 (2211) | 55665 (2742) |
| Multiplicity | 3.7 (3.6) | 13.5 (13.7) | 13.6 (14.2) |
| Completeness [%] | 96.6 (98.4) | 100.0 (100.0) | 100.0 (100.0) |
| Mean I/sigma(I) | 8.5 (1.5) | 9.6 (0.4) | 7.6 (0.1) |
| Wilson B-factor (Å<sup>2</sup>) | 33.4 | 43.0 | 39.0 |
| R<sub>meas</sub> [%] | 9.8 (82.7) | 18.3 (386.3) | 20.6 (503.4) |
| CC<sub>1/2</sub> | 0.996 (0.737) | 1.0 (0.300) | 1.0 (0.300) |
| Reflections used in refinement | 41939 (4195) | 44929 (4366) | 50418 (930) |
| R<sub>cryst</sub> / R<sub>free</sub> [%] | 20.7 (31.0) /<br>24.5 (33.7) | 21.1 (40.5) /<br>24.1 (46.7) | 22.6 (102.0) /<br>25.2 (126.2) |
| Total no. of non-hydrogen atoms (protein) | 3779 | 3810 | 3825 |
| No. of protein/ligand/solvent residues | 448 / 11 / 201 | 449 / 11 / 194 | 451 / 10 / 212 |
| RMSD bond length, bond angles [Å, °] | 0.011 / 1.09 | 0.012 / 1.08 | 0.011 / 1.21 |
| Ramachandran favoured/allowed/outliers/rotamer outliers [%] | 96.80 / 3.20 /<br>0.00 / 0.81 | 96.58 / 3.42 /<br>0.00 / 0.53 | 95.94 / 3.39 /<br>0.68 / 1.06 |
| Clashscore | 4.20 | 3.01 | 3.72 |
| Average B-factor/protein/ligands/solvent | 41.81 / 41.48 /<br>60.69 / 44.62 | 50.73 / 50.23 /<br>78.97 / 51.63 | 50.68 / 50.36 /<br>66.50 / 52.77 |

**Table S3.** Summary of calculated properties of nsp14 targeting fragment hits and analogues.
MW: molecular weight; cLogP: calculated octanol-water partition coefficient; HBA: hydrogen bond acceptor; HBD: hydrogen bond donor; tPSA: topological polar surface area.
MolLogS refers to water solubility.

| **Compound ID** | **MW [Da]** | **cLogP [mol]** | **HBA & HBD [No.]** | **tPSA [Å<sup>2</sup>]** | **MolLogS [Log(moles/L)] S [mg/L]** |
| --- | --- | --- | --- | --- | --- |
| **12A2** | 229.33 | 0.64 | 4 / 2 | 36.42 | -1.75*<br>4033.00 |
| **12D8** | 248.34 | 2.46 | 3 / 0 | 32.67 | -2.56<br>679.02 |
| **12F3** | 232.09 | 1.08 | 4 / 2 | 60.16 | -2.11<br>1799.83 |
| **12H4** | 214.27 | 1.99 | 2 / 0 | 32.67 | -2.03<br>1985.81 |
| **13A3** | 203.24 | 2.34 | 2 / 1 | 38.33 | -2.63<br>480.73 |
| **13C4** | 111.15 | -0.70 | 1 / 2 | 41.62 | -0.46<br>38468.70 |
| **3A23** | 161.20 | 1.32 | 1 / 2 | 32.26 | -1.26<br>8933.52 |
| **1D10** | 188.23 | 1.32 | 2 / 1 | 35.83 | -1.19<br>12091.69 |
| **1G7** | 162.14 | 2.45 | 3 / 1 | 46.53 | -2.31<br>789.88 |
| **6H5** | 157.19 | 0.31 | 3 / 2 | 60.16 | -1.58<br>4145.38 |
| **7B2** | 197.65 | 0.66 | 4 / 2 | 60.16 | -2.12<br>1492.65 |
| **12E3** | 226.07 | 1.61 | 3 / 2 | 60.16 | -2.53<br>668.07 |
| **Q11** | 246.12 | 1.58 | 4 / 2 | 60.16 | -2.18<br>1622.64 |
| **Q16** | 251.52 | 1.24 | 3 / 0 | 34.14 | -2.40<br>986.84 |
| **Q17** | 163.21 | -0.05 | 4 / 2 | 60.16 | -1.62<br>3891.56 |

\*First value in Log(moles/L), second value in mg/L.

**Table S4.** TSA results for full-length nsp14, ExoN and MTase domains in the absence and presence of fragments.

| Fragment ID | Full-length nsp14 |  |  | ExoN domain |  |  | MTase domain |  |  |
| --- | --- | --- | --- | --- | --- | --- | --- | --- | --- |
| | TSA<br>$T_i \pm \text{SD}$<br>[°C] | TSA<br>$\Delta T_i$ [°C] | TSA<br>$\vert \Delta T_i\vert $-3SD<br>[°C] | TSA<br>$T_i \pm \text{SD}$<br>[°C] | TSA<br>$\Delta T_i$ [°C] | TSA<br>$\vert \Delta T_i\vert $-3SD<br>[°C] | TSA<br>$T_i \pm \text{SD}$<br>[°C] | TSA<br>$\Delta T_i$ [°C] | TSA<br>$\vert \Delta T_i\vert $-3SD<br>[°C] |
| **nsp14** | 50.00 ± 0.00 | N/A | N/A | 47.37 ± 0.06 | N/A | N/A | 48.88 ± 0.04 | N/A | N/A |
| **nsp14 + 1% DMSO** | 49.90 ± 0.10 | N/A | N/A | 47.97 ± 0.06 | N/A | N/A | 48.55 ± 0.05 | N/A | N/A |
| ExoN 1 |  |  |  |  |  |  |  |  |  |
| **12A2** | 50.87 ± 0.12 | 0.97 | 0.61 | Atypical curve |  |  | 49.30 ± 0.07 | 0.75 | 0.54 |
| **12F3\*** | 49.57 ± 0.06 | -0.33 | 0.15 | 47.67 ± 0.25 | -0.30 | -0.45 | 48.03 ± 0.04 | -0.53 | 0.41 |
| ExoN 2 |  |  |  |  |  |  |  |  |  |
| **12D8** | 49.97 ± 0.06 | 0.07 | -0.11 | 45.83 ± 0.06 | -2.14 | 1.96 | 48.05 ± 0.09 | -0.50 | 0.23 |
| **12H4** | 49.93 ± 0.06 | 0.03 | -0.15 | 46.47 ± 0.12 | -1.50 | 1.14 | 48.25 ± 0.05 | -0.30 | 0.15 |
| **13C4** | 50.17 ± 0.06 | 0.27 | 0.09 | 46.77 ± 0.12 | -1.20 | 0.84 | 48.55 ± 0.05 | 0 | -0.15 |
| ExoN 3 |  |  |  |  |  |  |  |  |  |
| **13A3** | 49.87 ± 0.15 | -0.03 | -0.42 | 48.50 ± 0.17 | 0.53 | -1.04 | 48.05 ± 0.05 | -0.50 | 0.35 |

\*Also binds to binding sites Exon 2, MTase 1, MTase 2 and MTase 10.

**Table S5.** Number of commercially available fragment analogues and their hits.

| **Initial fragment hit** | **Number of purchased analogues** | **Number of analogue hits binding to nsp14** |
| --- | --- | --- |
| 13A3 | 42 | 2 |
| 12A2 | 4 | no hits |
| 12F3 | 11 | 6 |
| 12H4 | 11 | no hits |
| 13C4 | 12 | 1 |
| Obtained by fragment merging | 5 | no hits |

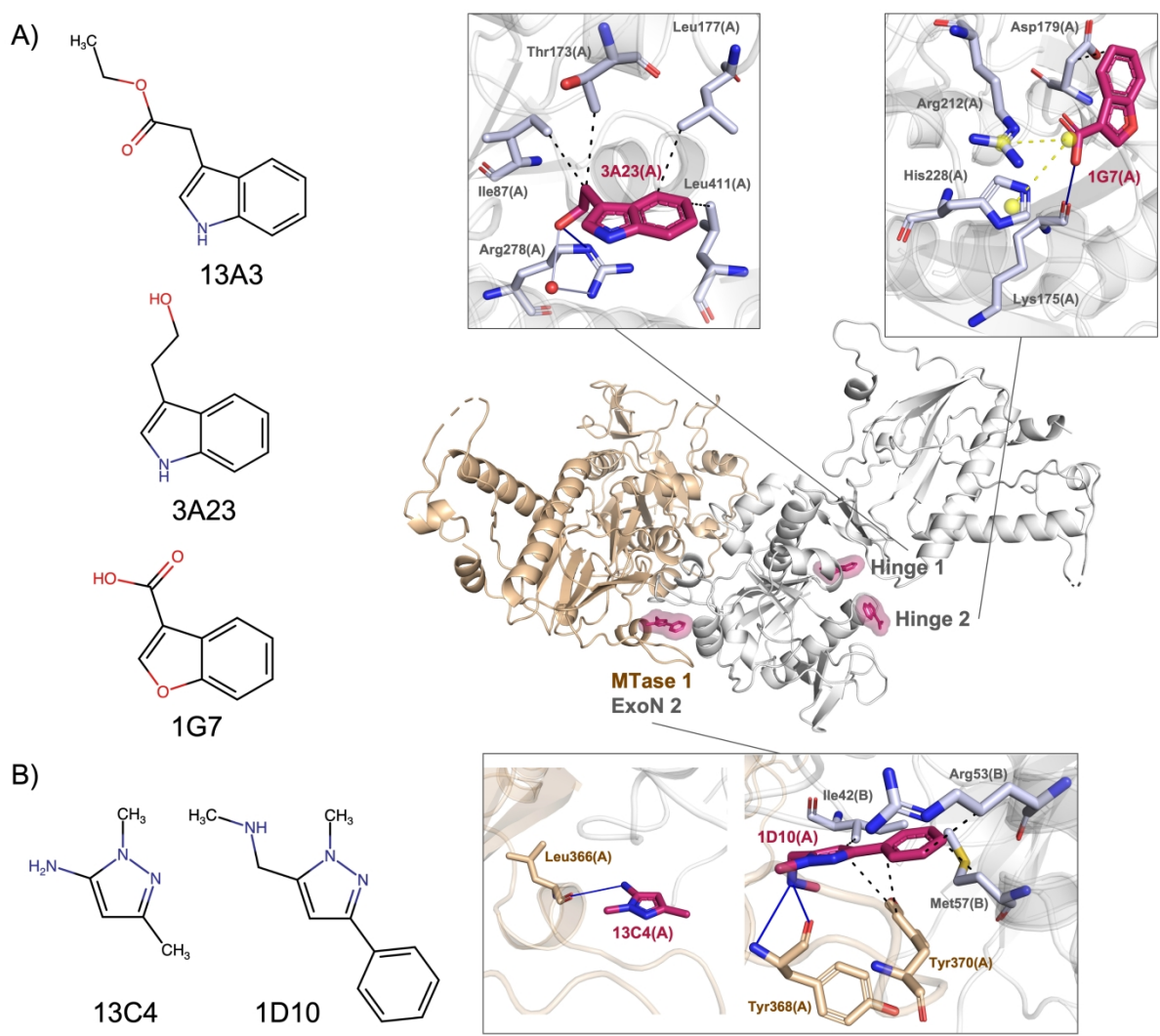

**Figure S1. Chemical structures and interactions of two fragments and their analogues with nsp14 residues.**

A) 13A3 and its analogues 3A23 and 1G7. B) 13C4 and its analogue 1D10. Nsp14 is coloured in grey. The relevant symmetry mate contributing to fragment binding is coloured in beige. The binding sites are shown in magenta surface. Hydrogen bonds are displayed as blue solid lines, hydrophobic interactions in black dashed lines, water bridges in grey solid lines and salt bridges in yellow dashed lines.

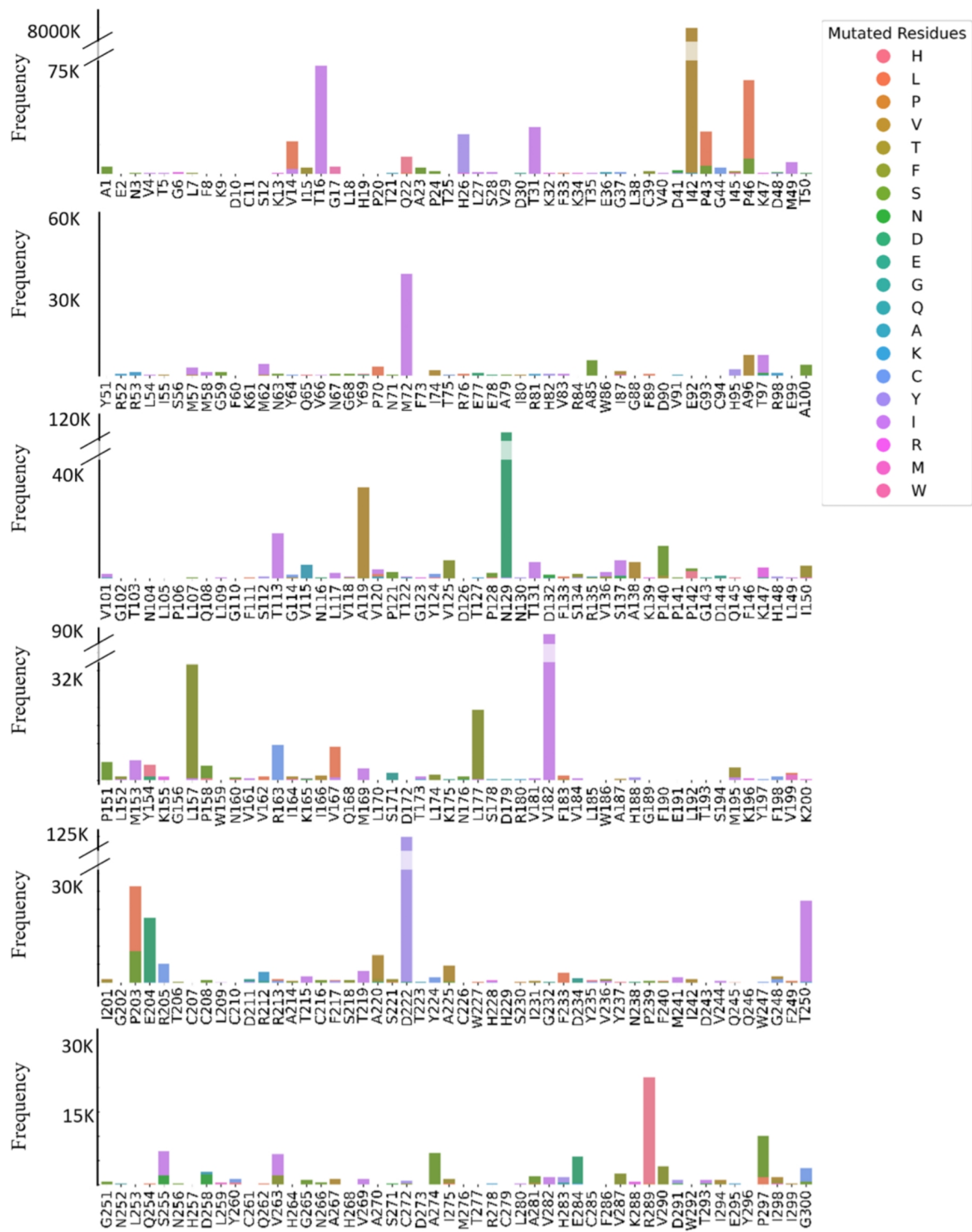

**Figure S2.** Mutation count of all 3435 mutations in nsp14 extracted from 14,246,785 nsp14 sequences, for residues 1 to 300.

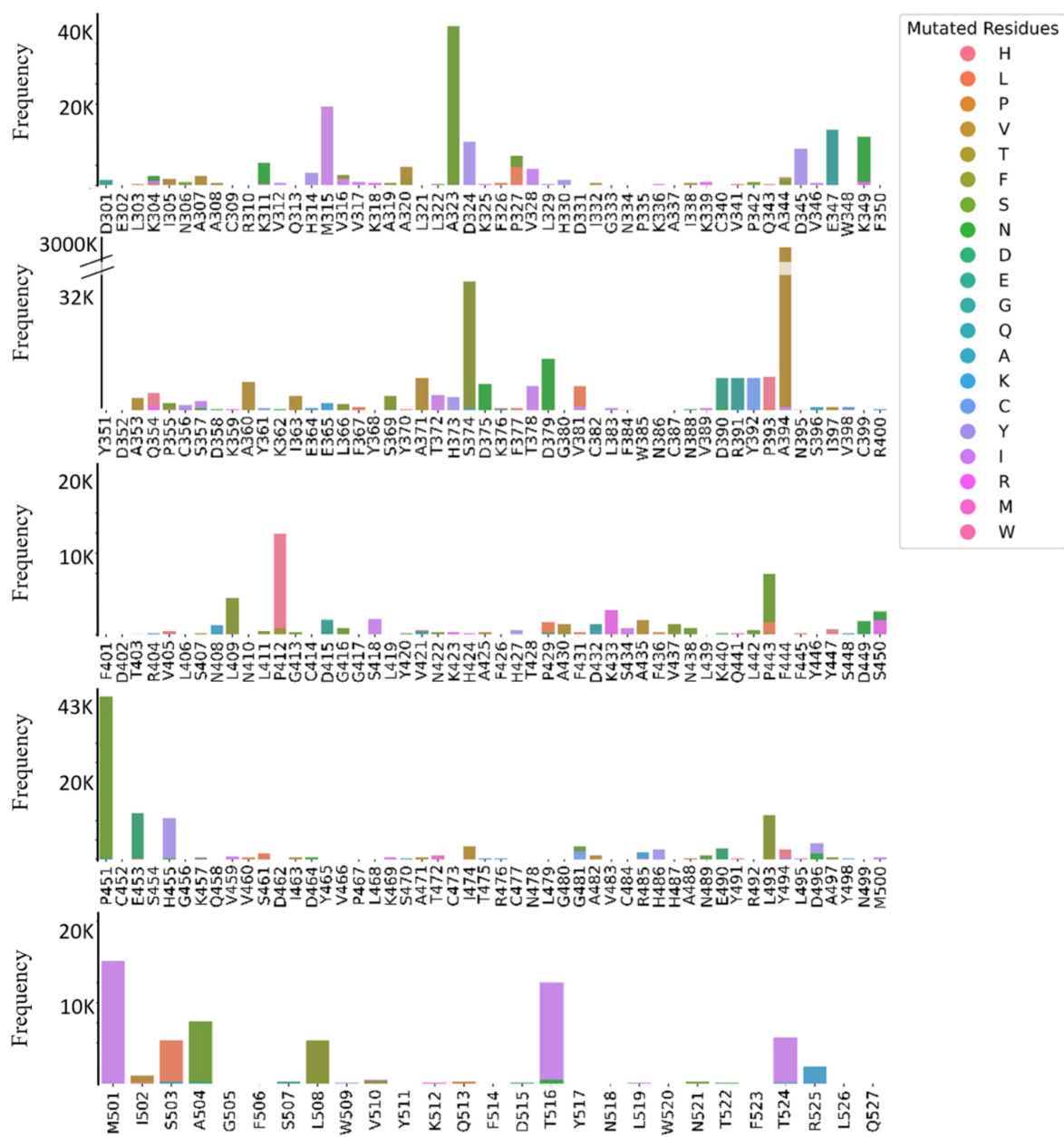

**Continuation Figure S2.** Mutation count of all 3435 mutations in nsp14 extracted from 14,246,785 nsp14 sequences, arranged for residues 301 to 527.

**Table S6.** Summary of deposited coordinates of nsp14 targeting fragments, their domain binding regions, description of binding pockets and suggested naming system for the binding sites. Our novel fragment hits are coloured in blue, previously reported fragment hits are coloured in black (Imprachim *et al.*, 2023; Meyer *et al.*, 2025). Compounds binding to pockets formed by one nsp14 and its symmetry mate are listed in both binding sites.

| PDB | Binding region(s) | Binding pocket | Name of binding site |
| --- | --- | --- | --- |
| 5SM2, **12F3**, **12A2**, 6H5, 7B2, Q17 | ExoN | nsp10 interface | ExoN 1 |
| 5SME, 5SMF, 5SLZ, 5SLU, 5SLE, 5SLV, 5SLX, 5SMI, **12F3**, **12D8**, **12H4**, **13C4**, 6H5, 7B2, Q11, 12E3, 1D10, Q17 |  |  | ExoN 2 |
| 5SLR, **13A3** |  |  | ExoN 3 |
| 5SLL, 5SKX |  |  | ExoN 4 |
| 5SLU, 5SLF, 5SMF (2<sup>nd</sup>) |  |  | ExoN 5 |
| 5SKY, 5SMD |  | ExoN active | ExoN 6 |
| **12E3** |  | ExoN other | ExoN 7 |
| 5KSW, 5SKZ, 5SL4, 5SLH, 5SLW, 5SM6, **3A23** | Hinge (and ExoN) | Hinge region | Hinge 1 |
| 5SL3, 5SL5, 5SLS, 5SLT, **1G7** |  | Between ExoN and MTase | Hinge 2 |
| 5SL0, 5SL1, 5SL2, 5SL6, 5SLI, 5SLJ, 5SLM, 5SLN, 5SLX, 5SLY, 5SM4, 5SM9, 5SMA, 5SMC, **12F3**, **12D8**, **12H4**, **13C4**, 6H5, 7B2, Q11, 12E3, 1D10, Q17 | MTase | MTase allosteric | MTase 1 |
| 5SL9, 5SLA, 5SLB, 5SLC, 5SLD, 5SLE (2<sup>nd</sup>), 5SLF (2<sup>nd</sup>), 5SLG, 5SLK, 5SLO, 5SLW (2<sup>nd</sup>), 5SM0, 5SM2 (2<sup>nd</sup>), 5SM7, 5SM8, 5SMB, 5SMC (2<sup>nd</sup>), 5SMG, **12F3**, **Q16**, TDI-015051 |  | MTase active | MTase 2 |
| 5SL6 (2<sup>nd</sup>), 5SM3, 5SMH |  | MTase other | MTase 3 |
| 5SLA (2<sup>nd</sup>), **13A3** |  |  | MTase 4 |
| 5SL4 (2<sup>nd</sup>), 5SM5 |  |  | MTase 5 |
| 5SL7, 5SLQ, 5SM1 |  |  | MTase 6 |
| 5SL8 |  |  | MTase 7 |
| 5SLI (2<sup>nd</sup>), **12E3** |  |  | MTase 8 |
| 5SLP |  |  | MTase 9 |
| 5SL0 (2<sup>nd</sup>), 5SLI (3<sup>rd</sup>), **12F3**, 6H5 |  |  | MTase 10 |

**Table S7:** Ranking of fragment binding sites on SARS-CoV-2 nsp14. <sup>\*1</sup>Residues with low mutation frequency are coloured in green, whereas those shaded in red show a high mutation frequency. <sup>\*2</sup>Using MERS, SARS and SARS-CoV-2 nsp14 protein sequences. <sup>\*3</sup>Ranked first as proof-of-principle data are available (Meyer *et al.*, 2025). <sup>\*4</sup>Fragments binding to this site may disrupt the conformational rearrangement of the SAM-interacting loop by stabilising an alternative structural state, thereby preventing it from adopting the conformation observed in SAM-bound nsp10–nsp14 complexes (Imprachim *et al.*, 2023).

| Rank | Binding sites Suggested names | Function / Validation | <sup>*1</sup>Mutation in binding site? | <sup>*2</sup>Protein Sequence alignment High / low conservation | Binding site formed with a symmetry mate? | Biophysical evaluation? | Number of bound fragments per site | PAINS |
| --- | --- | --- | --- | --- | --- | --- | --- | --- |
| 1 | <sup>*3</sup>MTase 2 | MTase active site |  | high | No | Yes | 20 |  |
| 2 | ExoN 6 | ExoN active site | E92, P106, H268, D273 | high | No | No | 2 |  |
| 3 | MTase 4 | close to nsp12 interface | N478 | high | Yes | No | 2 |  |
| 4 | ExoN 4 | nsp10 interface |  | low | No | No | 2 |  |
| 5 | MTase 9 | close to nsp12 interface |  | low | No | No | 1 |  |
| 6 | ExoN 7 | nsp7 interface |  | low | Yes | Yes | 1 |  |
| 7 | ExoN 3 | close to nsp10 interface |  | low | Yes | Yes | 2 |  |
| 8 | ExoN 1 | nsp10 interface | M72, G202 | high | No | Yes | 6 |  |
| 9 | ExoN 5 | nsp10 interface | P106, A119 | high | No | No | 3 |  |
| 10 | Hinge 1 | Hinge region | W86, T113, L177 | high | No | Yes | 7 | 5SKZ |
| 11 | Hinge 2 | Hinge region | L117 | low | No | No | 5 |  |
| 12 | MTase 1 | <sup>*4</sup>MTase allosteric | S374 | low | Yes | Yes | 24 | 5SLX |
| 13 | ExoN 2 | nsp10 interface | I42, P43, P46 | low | Yes | Yes | 18 | 5SLX |
| 14 | MTase 5 | MTase other | R310 | low | No | No | 2 |  |
| 15 | MTase 3 | MTase other |  | low | No | No | 3 |  |
| 16 | MTase 8 | MTase other |  | low | Yes | No | 2 |  |
| 17 | MTase 6 | MTase other |  | low | No | No | 3 |  |
| 18 | MTase 7 | MTase other | A323, N518 | low | No | No | 1 |  |
| 19 | MTase 10 | MTase other | A394 | low | No | No | 4 |  |

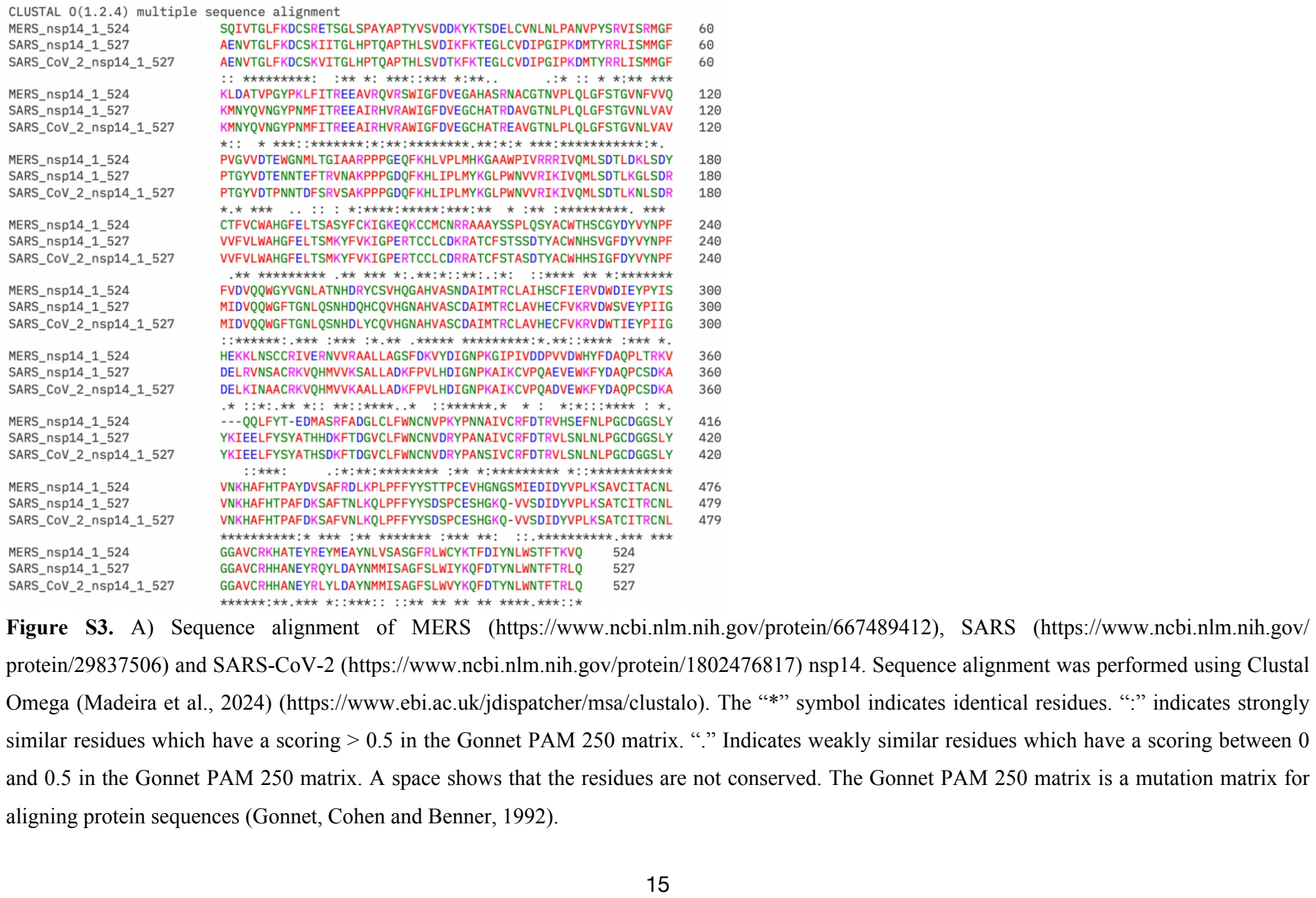

Figure S3. A) Sequence alignment of MERS (https://www.ncbi.nlm.nih.gov/protein/667489412), SARS (https://www.ncbi.nlm.nih.gov/ protein/29837506) and SARS-CoV-2 (https://www.ncbi.nlm.nih.gov/protein/1802476817) nsp14. Sequence alignment was performed using Clustal Omega (Madeira et al., 2024) (https://www.ebi.ac.uk/jdispatcher/msa/clustalo). The “\*” symbol indicates identical residues. “:” indicates strongly similar residues which have a scoring > 0.5 in the Gonnet PAM 250 matrix. “.” Indicates weakly similar residues which have a scoring between 0 and 0.5 in the Gonnet PAM 250 matrix. A space shows that the residues are not conserved. The Gonnet PAM 250 matrix is a mutation matrix for aligning protein sequences (Gonnet, Cohen and Benner, 1992).

**Table S8:** Conserved nsp14 residues in binding sites are shown in different colours according to the alignment results in Figure S3. Identical residues are coloured in red, strongly similar residues are shaded in orange, weakly similar residues are shown in yellow and non-conserved residues are shown in black.

| Binding sites | Residues in the binding sites |
| --- | --- |
| ExoN 1 | V29, R52, I55, F60, K61, M62, N63, Y64, Y69, P70, N71, M72, I201, G202, N238, P239, F240 |
| ExoN 2 | L38, I42, P43, G44, I45, P46, M49, R53, L54, M57, M58, T193 |
| ExoN 3 | N160, I164, T219, A220, G232, D234 |
| ExoN 4 | G59, F60, K61, K196, V199, K200, I201, F217, T219, Y235 |
| ExoN 5 | A96, T97, T103, N104, L105, P106, L107, A119, V120, P121, W159, F190, T193, S194, Y197 |
| ExoN 6 | D90, E92, G93, C94, P106, L107, Q108, V118, N266, H268, V269, A270, D273 |
| ExoN 7 | I164, K165, Q168 |
| Hinge 1 | W86, I87, S112, T113, T173, L177, R278, L406, N410, L411, P412 |
| Hinge 2 | L177, S178, D179, R212, H228, I299 |
| MTase 1 | Y351, E365, L366, F367, Y368, S369, T372, S374, D375, F377 |
| MTase 2 | N306, G333, N334, P335, A353, Q354, P355, C356, W385, N386, C387, N388, F401, Y420, N422, F426, T428, F506 |
| MTase 3 | E295, Y296, P297, I298, E302, L303, N306, N422, K423 |
| MTase 4 | K469, S470, A471, N478, L479, W520 |
| MTase 5 | R310, H314, I338, K339, C340, P342 |
| MTase 6 | K318, L321, L322, K325, F326, P327, V341, Q343, A344, D345, V346 |
| MTase 7 | A323, D324, Q513, N518 |
| MTase 8 | A471, T472, I474, T516, Y517, W520 |
| MTase 9 | V459, V460, D464, Y465, V466, R485, A488, N489, R492, L493 |
| MTase 10 | D324, F326, G380, A394, N395, Y511, Q513 |

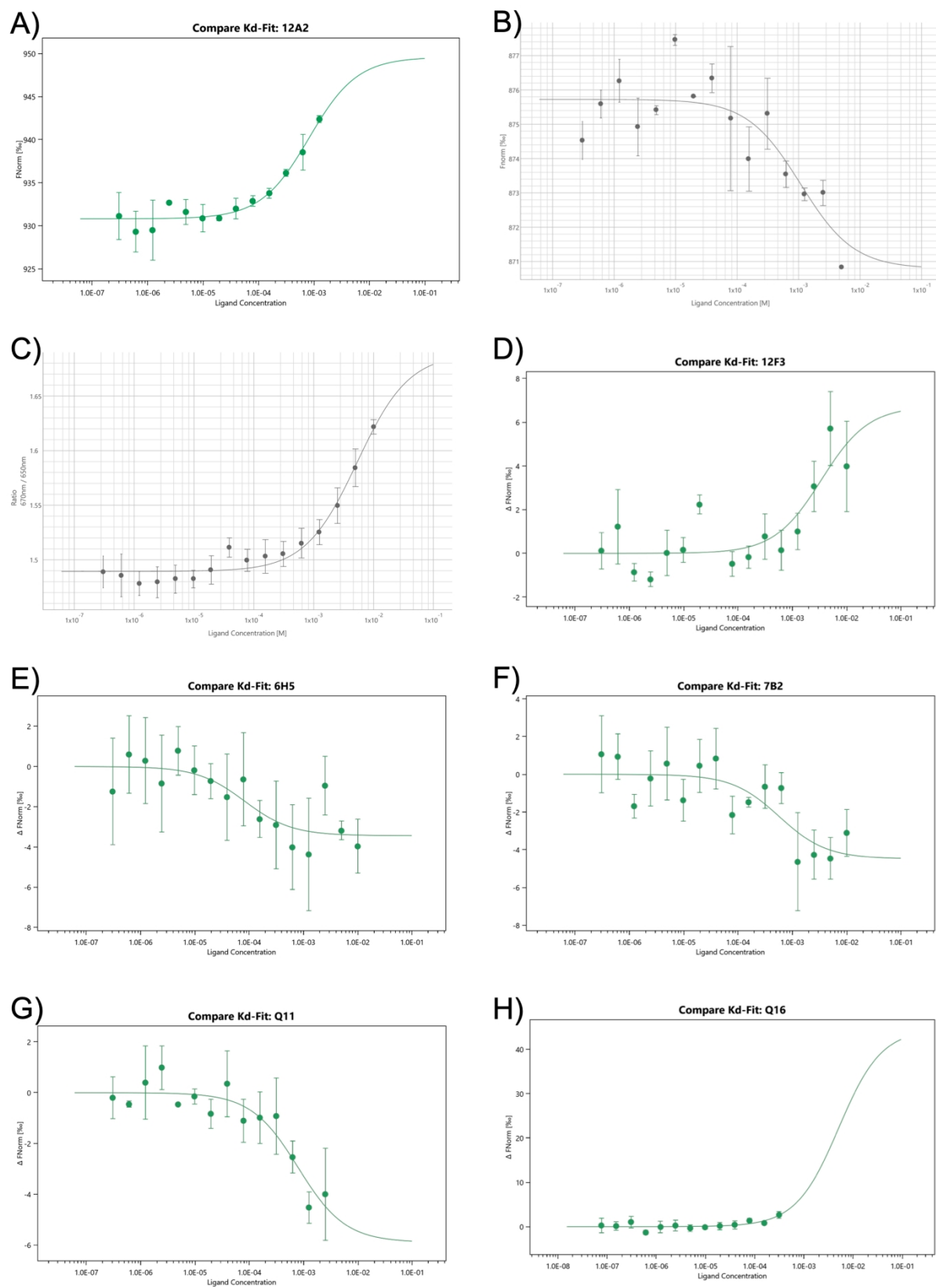

**Figure S4.** Binding curves for the binding interactions between fragments (analogues) and nsp14 in MST assays.

A) 12A2. B) 12H4. C) 12D8. D) 12F3. E) 6H5. F) 7B2. G) Q11. H) Q16.

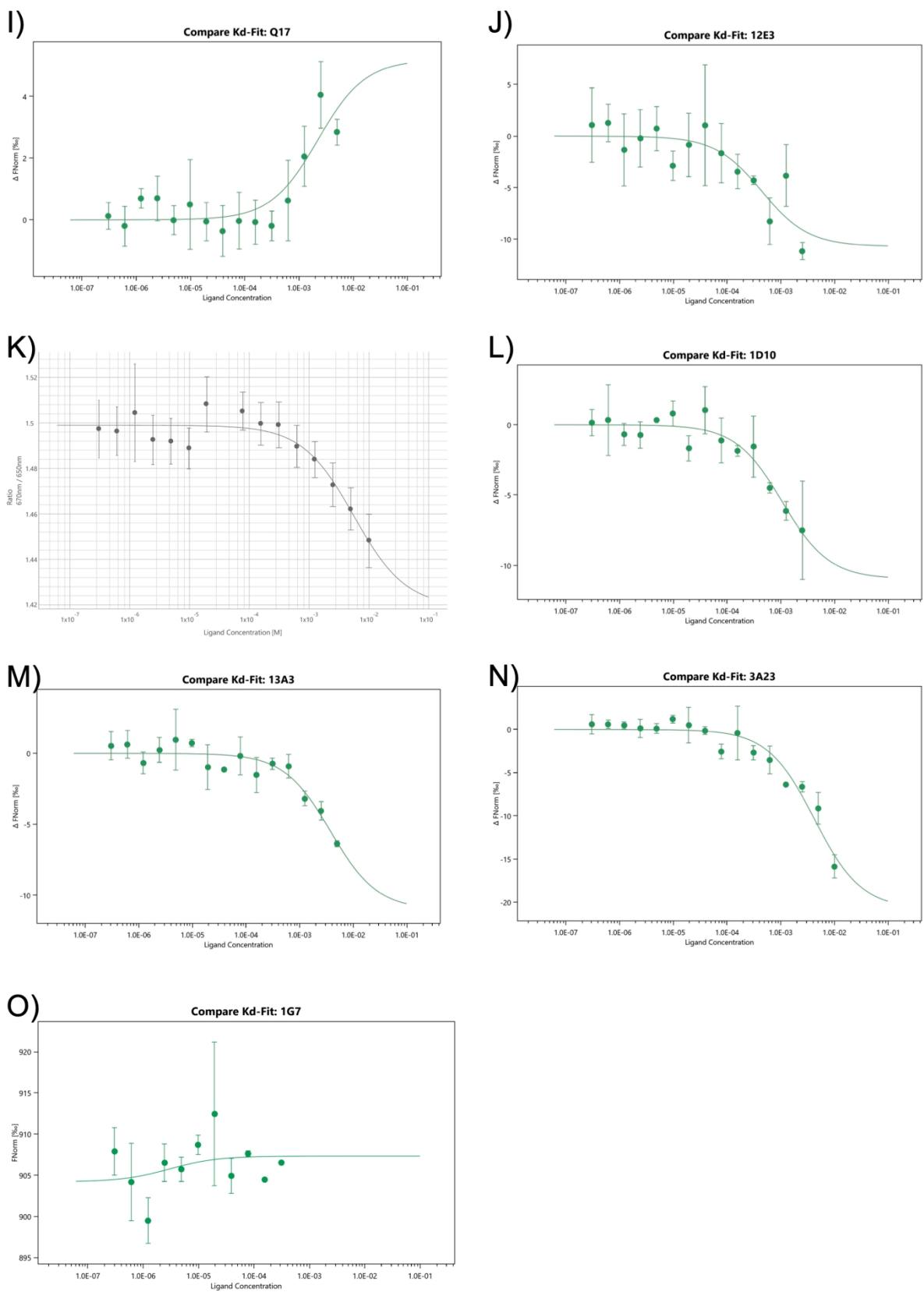

**Continuation Figure S4.** Binding curves for the binding interactions between fragments (analogues) and nsp14 in MST assays.

I) Q17. J) 12E3. K) 13C4. L) 1D10. M) 13A3. N) 3A23. O) 1G7.

### References (Supplementary Material)

Gonnet, G. H., Cohen, M. A. and Benner, S. A. (1992). Exhaustive Matching of the Entire Protein Sequence Database. *Science* **256**, 1443–1445.

Imprachim, N., Yosaatmadja, Y. and Newman, J.A. (2023). Crystal structures and fragment screening of SARS-CoV-2 NSP14 reveal details of exoribonuclease activation and mRNA capping and provide starting points for antiviral drug development. *Nucleic Acids Res.* **51**, 475–487.

Madeira, F., Madhusoodanan, N., Lee, J., Eusebi, A., Niewielska, A., Tivey, A. R. N., Lopez, R. and Butcher, S. (2024). The EMBL-EBI Job Dispatcher sequence analysis tools framework in 2024. *Nucleic Acids Res.* **52**(W1), W521–W525.

Meyer, C., Garzia, A., Miller, M. W., Huggins, D. J., Myers, R. W., Hoffmann, H. H., Ashbrook, A. W., Jannath, S. Y., Liverton, N., Kargman, S., Zimmerman, M., Nelson, A. M., Sharma, V., Dolgov, E., Cangialosi, J., Penalva-Lopez, S., Alvarez, N., Chang, C. W., Oswal, N., Gonzalez, I., ... Tuschl, T. (2025). Small-molecule inhibition of SARS-CoV-2 NSP14 RNA cap methyltransferase. *Nature* **637**(8048), 1178–1185.
